# piRNA-mediated gene regulation and adaptation to sex-specific transposon expression in *D. melanogaster* male germline

**DOI:** 10.1101/2020.08.25.266585

**Authors:** Peiwei Chen, Alexei A. Kotov, Baira K. Godneeva, Sergei S. Bazylev, Ludmila V. Olenina, Alexei A. Aravin

## Abstract

Small non-coding piRNAs act as sequence-specific guides to repress complementary targets in Metazoa. Prior studies in *Drosophila* ovaries have demonstrated the function of piRNA pathway in transposon silencing and therefore genome defense. However, the ability of piRNA program to respond to different transposon landscape and the role of piRNAs in regulating host gene expression remain poorly understood. Here, we comprehensively analyzed piRNA expression and defined the repertoire of their targets in *Drosophila melanogaster* testes. Comparison of piRNA programs between sexes revealed sexual dimorphism in piRNA programs that parallel sex-specific transposon expression. Using a novel bioinformatic pipeline, we identified new piRNA clusters and established complex satellites as dual-strand piRNA clusters. While sharing most piRNA clusters, two sexes employ them differentially to combat sex-specific transposon landscape. We found several host genes targeted by piRNAs in testis, including *CG12717/pita*, a SUMO protease gene. piRNAs encoded on Y chromosome silence *pita*, but not its paralog, to exert sex- and paralog-specific gene regulation. Interestingly, *pita* is targeted by endogenous siRNAs in a sibling species, *Drosophila mauritiana*, suggesting distinct but related silencing strategies invented in recent evolution to regulate a conserved protein-encoding gene.

## INTRODUCTION

PIWI-interacting (pi)RNA is a class of small non-coding RNAs named after their interaction with PIWI-clade Argounate proteins. piRNAs guide PIWI proteins to complementary RNAs, thereby specifying the target of PIWI silencing. Unlike miRNAs and siRNAs that are ubiquitously expressed, the expression of piRNAs is restricted to gonads in many animals. As a result, perturbation of the piRNA program often compromises reproductive functions with no obvious defects in soma. *Drosophila melanogaster* is one of the most used model organisms to study piRNA biogenesis and function. In fact, piRNAs were first described in fly testes (Aravin et al., 2001; Vagin et al., 2006). However, most subsequent studies were performed using ovaries as a model system. Work on female gonads has shown that most piRNAs have homology to transposable elements (TEs), suggesting TEs as major targets of piRNAs (Brennecke et al., 2007). Studies on fly ovaries also identified large intergenic regions dubbed piRNA clusters that harbor nested TE fragments, which act as genomic source loci of piRNAs. A peri-centromeric region on chr2R called *42AB* was found to be the most active piRNA cluster in ovaries. It remains largely unexplored to what extent these findings from ovaries are applicable to the male counterpart. To date, we still know very little about how sexually dimorphic the *Drosophila* piRNA program is, besides that there is a single locus on Y chromosome called *Suppressor of Stellate (Su(Ste)*) that produces piRNAs only in males.

Importantly, *Drosophila* as an animal model offers unique value to studying sexual dimorphism of the piRNA program in general. In zebrafish, piRNA pathway mutants are always phenotypically males (Houwing et al., 2007, 2008; Kamminga et al., 2010), rendering it nearly impossible to probe the impact of piRNA loss in females. In mice, an intact piRNA program is only required for male fertility, while murine females are insensitive to piRNA loss (Carmell et al., 2007; Deng and Lin, 2002; Kuramochi-Miyagawa et al., 2004). Contrary to fish and mouse, fly fertility is dependent on a functional piRNA pathway in both sexes (Aravin et al., 2001; Brennecke et al., 2007; Lin and Spradling, 1997; Vagin et al., 2006). Therefore, *Drosophila* provides an unparalleled opportunity to study whether, and if so how, the piRNA program can be modified in each sex to safeguard reproductive functions.

In this study, we comprehensively analyzed the piRNA profile in *Drosophila melanogaster* testis and compared it to the female counterpart. Besides TEs, we found complex satellites as another class of selfish genetic elements targeted by the piRNA pathway in gonads of both sexes. Our analysis showed that TE-silencing piRNA program is sexually dimorphic, and it shows evidence of adaptation to sex-specific TE landscape. To understand the genomic origins of differentially produced piRNAs, we sought to *de novo* define genome-wide piRNA clusters in testis. However, we noticed that the standard pipeline used for ovary piRNAs failed to detect known piRNA clusters in testis, so we developed a new bioinformatic algorithm to tackle this problem. Using the new algorithm, we were able to identify novel piRNA clusters and to quantify their activities more accurately in both sexes. Notably, piRNA source loci are employed differentially in males and females, and the sex bias of piRNA cluster expression appears to match that of their TE contents. We also found two loci producing piRNAs with the potential to repress host protein-encoding genes, including a newly identified locus on Y that produces piRNAs against *CG12717/pita*. Expression of *pita*, but not its paralog *veloren*, is de-repressed in multiple piRNA pathway mutants, indicating that piRNAs silence its expression and can distinguish paralogs with sequence similarities. Finally, we explored the evolutionary history of *pita* and found it to be a young gene conserved in the *melanogaster* subgroup. Intriguingly, *pita* is targeted by another class of small non-coding RNAs, endogenous siRNAs, in the sibling species *Drosophila mauritiana*, suggesting distinct small RNA-based silencing strategies invented in recent evolution to regulate a young yet conserved gene.

## RESULTS

### *Drosophila* piRNA program is sexually dimorphic

To characterize the piRNA profile in male gonads, we sequenced 18-30nt small RNAs from testes and compared them with published ovary small RNA datasets (ElMaghraby et al., 2019). Mapping and annotation of small RNA reads using the pipeline shown in Figure S1 revealed large differences in the expression of major classes of small RNAs between testes and ovaries. In agreement with previous findings (Czech et al., 2008), TE-mapping 23-29nt piRNAs are the most abundant class of small RNAs in ovaries, while 21-23nt microRNAs constitute a minor fraction and an even smaller one for 21nt endogenous (endo-) siRNAs (Figure 1A). In contrast, miRNAs constitute a larger fraction in testes, so do endo-siRNAs that map to protein-encoding genes, consistent with a previous report (Wen et al., 2015). To define the piRNA population, we eliminated reads mapping to other types of non-coding RNA (rRNA, miRNA, snRNA, snoRNA and tRNA) from 23-29nt small RNAs. Remaining reads show a strong bias for U at the first nucleotide (“1U bias”: 70.9%), the feature of *bona fide* piRNAs (Figure 1B). The piRNA-to-miRNA ratio is distinct between sexes: ~10 in ovary and ~2 in testis. In both sexes, piRNAs mapping to TEs take up the largest fraction of total piRNAs. However, whereas 66% of piRNAs mapped to TEs in ovaries, only 40% mapped to TEs in testes (Figure 1B). Meanwhile, larger fractions of total piRNAs mapped to protein-encoding genes (including introns) and intergenic regions in testes (24.6% and 30.0%, respectively) than ovaries (19.6% and 10.7%, respectively). These results suggest that distinct piRNA programs operate in male and female gonads.

**Figure 1.**
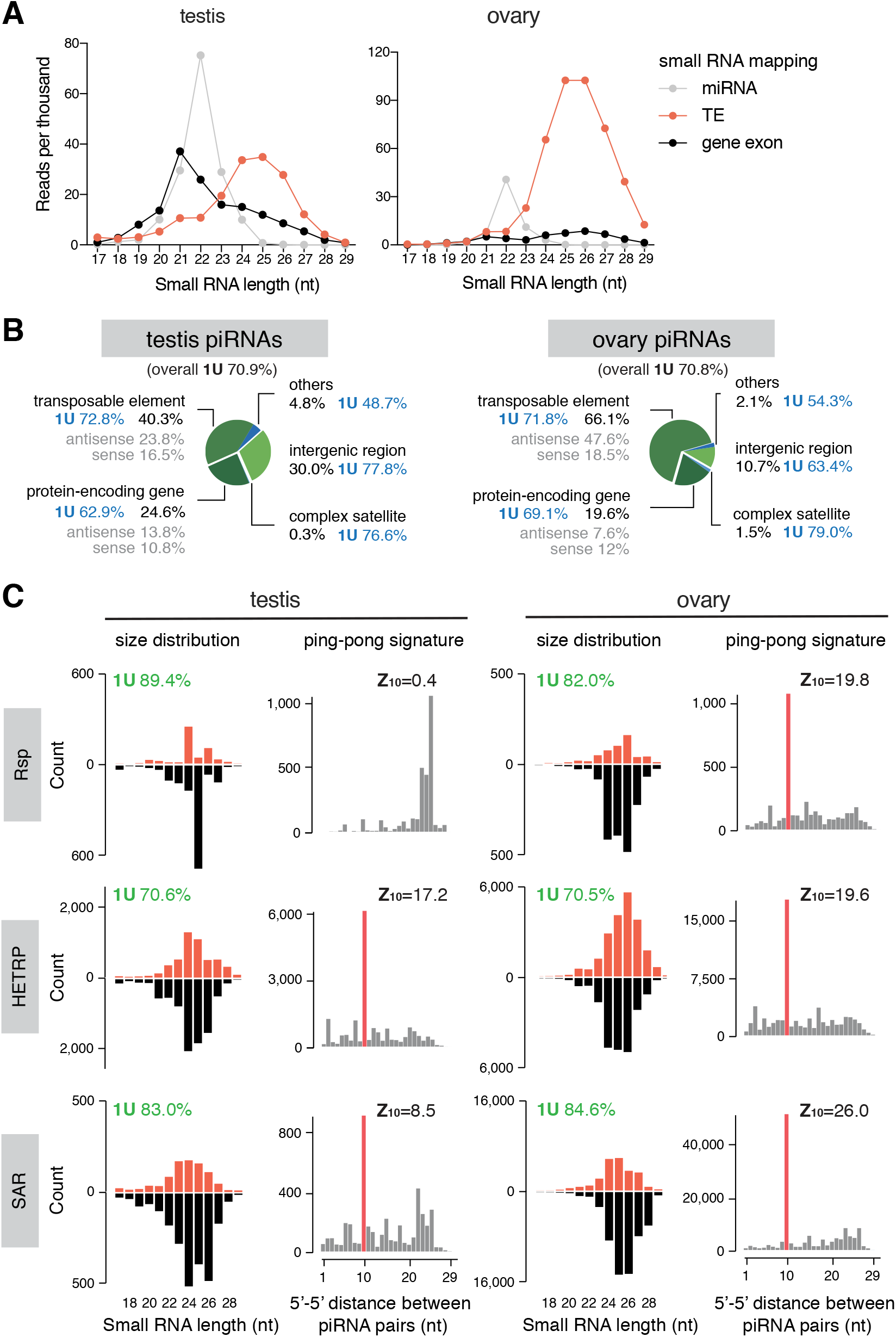
Analysis of small RNA profiles in testis and ovary. (A) Size distribution plots of microRNAs (gray), remaining small RNAs that map to TE consensus (red) and protein-encoding gene exons (black), in testis (left) and ovary (right). (B) Annotation of piRNA reads in testis (left) and ovary (right). 1U nucleotide bias (%) for overall piRNA population and each category is shown next to labels. See also Figure S1. (B) Characterization of piRNAs mapping to three known complex satellites in two sexes. Left panels of each sex are size distribution of piRNAs mapping to consensus sequences of each complex satellite. Right panels are distributions of 5’-to-5’ distances of piRNA pairs, showing an enrichment for 10nt (i.e., ping-pong signature), except for *Rsp* in testis. 1U nucleotide bias (%) and ping-pong z-score are shown above plots. See also Figure S2.

Testis piRNAs also map to several known complex satellites: *HETRP/TAS* (a sub-telomeric satellite repeat), *Responder (Rsp)* and *SAR* (related to *1.688* repeat family) (Figure 1C; Figure S2A). Complex satellite-mapping small RNAs in testis exhibit 1U bias and size distribution that peaks around 24-26nt, consistent with their piRNA identities. Both strands of complex satellites produce piRNAs, and their production depends on Rhi (see accompanying manuscript), a protein that marks dual-strand piRNA clusters and is required for their expression (Klattenhoff et al., 2009; Mohn et al., 2014; Zhang et al., 2014). Similarly, ovary small RNAs also map to complex satellites and show features of *bone fide* piRNAs, including 1U bias, size distribution that peaks around 24-26nt, small RNA production from both strands and dependency on Rhino. Moreover, piRNAs from complex satellites show ping-pong signature, an enrichment for 10nt overlap between the 5’ ends of complementary piRNA pairs, except for *Rsp* in testis (Figure 1C; Figure S2B). These results show that complex satellites are sources of piRNAs in both sexes, pointing to a possible role of piRNAs in regulating satellite DNA and associated heterochromatin in the gonad.

We next analyzed piRNAs targeting different TE families. Comparison of small RNA profiles in testis and ovary showed that piRNAs targeting different TEs are expressed at different levels in two sexes (Figure 2A). Top 3 TEs targeted by piRNA are all different in testis and ovary, and, among top 10, only 3 are shared between sexes (Figure 2B). The most differentially targeted TEs are two telomere-associated TEs, *HeT-A* and *TAHRE*, which ovary makes 106 and 74 times more antisense piRNAs, respectively, than testis. In contrast, several elements such as *baggins1, invader3* and *copia* are targeted by more piRNAs in testis. piRNAs targeting all but one (*copia*) TE families show stronger ping-pong signature in ovary, as measured by ping-pong z-score (Figure 2A). In conclusion, different TE families are targeted by piRNAs differentially in two sexes.

**Figure 2.**
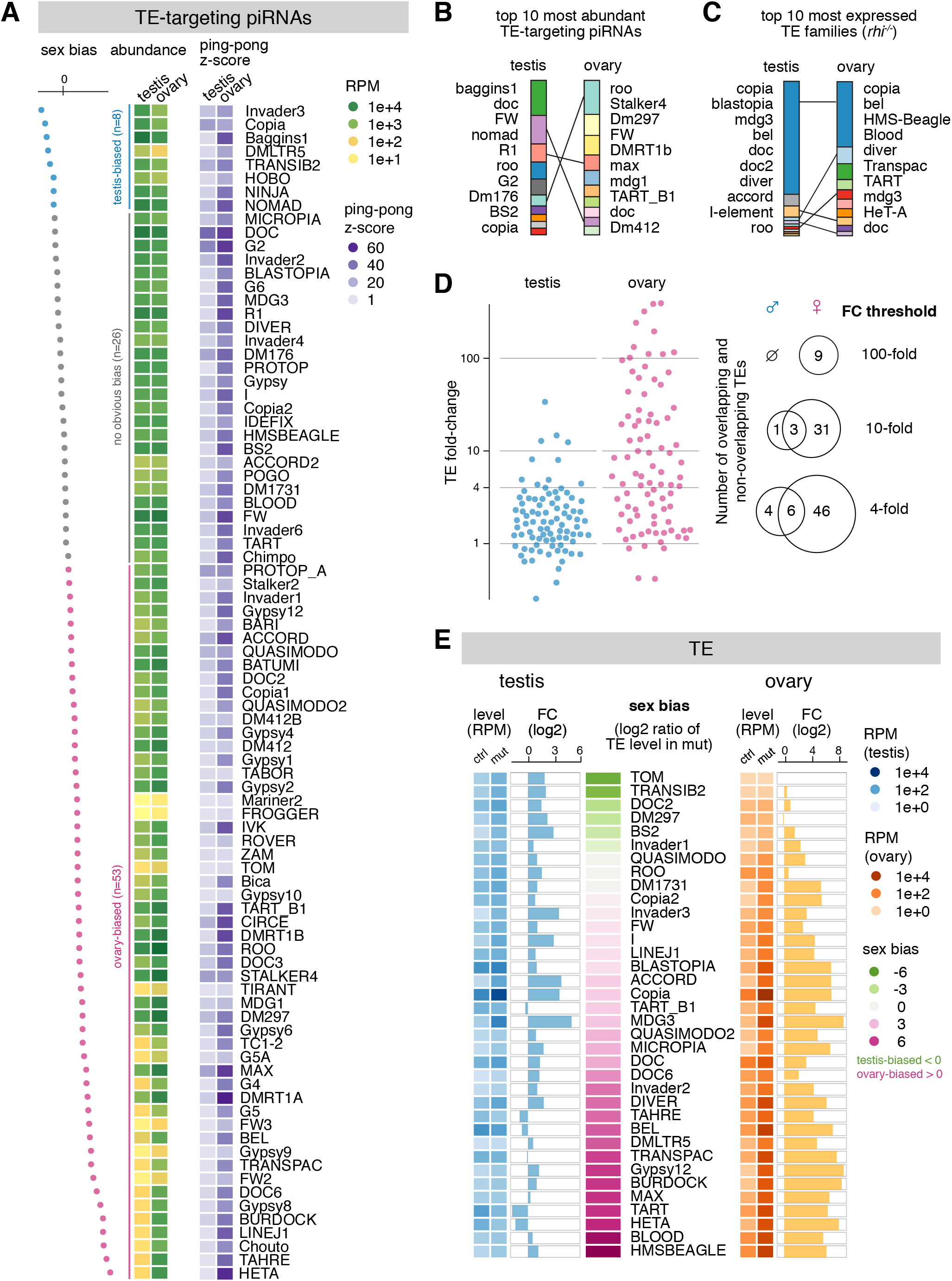
Expression of piRNAs and TEs are both sexually dimorphic. (A) Heatmaps showing the abundance of antisense piRNA (left) and ping-pong z-score (right) for each TE family in two sexes. TE families are sorted by sex bias of piRNA expression, defined as the log2 ratio of antisense piRNA abundance in testis over ovary. TEs with more than 2-fold differences in antisense piRNAs are colored as testis-biased (blue) and ovary-biased (pink), respectively, with the remaining having no obvious bias (gray). (B) Top 10 TEs targeted by the most abundant antisense piRNAs in testis (left) and ovary (right). Heights of slices correspond to relative abundance in each sex, and the sum of top 10 TEs is then scaled to the same height between sexes. Each TE family is given a unique color, and the same TE family is connected by a line to help visualize distinct rank-orders between sexes. Names of TE families are shown following the same order, though not directly next to respective slices. (C) Top 10 most expressed TE families in piRNA pathway mutant testis (left) and ovary (right). *rhi^-/-^* was used, where piRNA production from genome-wide dual-strand piRNA clusters collapses. Slice heights and colors were depicted as described in (B), though the same TE can be marked by a different color from (B). (D) Scatter plot displaying the fold-change of TE expression in piRNA pathway mutant (*rhi^-/-^*) testis (left, blue) and ovary (right, pink) over controls. Venn diagrams of the number of TEs showing 100-, 10- and 4-fold de-repression in two sexes are shown on the right. (E) Expression of 36 TE families that are regulated by *rhi* (see methods) in testis (left) and ovary (right). TE families are sorted by sex bias of their expression in piRNA pathway mutant (*rhi^-/-^*), defined as the log2 ratio between sexes. Heatmaps display TE levels in control and mutant, while bar graphs show the fold-change of expression in mutant over control.

### Distinct piRNA programs in two sexes parallel sex-specific TE expression

To explore if sex differences in TE-targeting piRNA programs are accompanied by differential expression of TEs themselves, we set out to compare expression levels of different TE families in two sexes. Since piRNA pathway efficiently represses TEs, their expression in wild-type animals does not reflect their full expression potentials that can be achieved when piRNA silencing is removed. Hence, we analyzed TE expression in testes and ovaries of *rhi* mutants that lose piRNA production from dual-strand clusters in both sexes (see accompanying manuscript) and controls.

Profiling TE expression in two sexes by polyA-selected (polyA+) RNA-seq demonstrated clear sexual dimorphism. Overall, TE expression in piRNA pathway mutant testes and ovaries is weakly correlated (Spearman’s *ρ*: 0.26; Figure S3A). Among the 10 most expressed TE families in two sexes, only 5 overlap, though the same element, *copia*, has the highest expression in both ovary and testis (Figure 2C). There are more TE families expressed above each of the three expression cutoffs (1000, 100 and 25 RPM) in ovaries than testes (Figure S3A). The most ovary-biased TEs include *Blood, max, Burdock* and two telomere-associated TEs, *HeT-A* and *TART* (Figure 2E). Only a few TE families are expressed higher in testis than ovary (Figure 2E; Figure S3A). In this group, *doc2* shows the highest expression in testis (87 RPM, 8.5-fold higher than ovary). Several elements are expressed at lower levels but have stronger biases for expression in testis: expression of *Tom* could only be detected in testis but not in ovary, and *Transib2* is expressed 28-fold higher in testis than ovary. Overall, the majority of TE families demonstrate strong differences in their expression between sexes.

To quantify the effect of piRNA pathway in suppressing TEs in two sexes, we calculated levels of TE de-repression upon disruption of piRNA pathway. Few TE families remained unaffected by *rhi* mutation, often accompanied by unperturbed antisense piRNA production (e.g., *gypsy, gypsy10* and *tabor*). There are 9 TE families up-regulated more than 100-fold in ovary. In contrast, no TE is up-regulated that strongly in testis (Figure 2D). Overall, the vast majority of TEs show stronger de-repression in ovaries, with *gypsy12* (389-fold), *Burdock* (317-fold), *HeT-A* (239-fold) and *TART* (80-fold) being the most prominent examples, as all of them exhibited no or mild de-repression (<4-fold) in testes (Figure 2E). We found only 6 TEs that show stronger (at least 4-fold) de-repression in testis than ovary (*Transib2, BS2, baggins1, Dm297, invader3, invader6*). Altogether, our results show that piRNAs regulate the expression of different TE families to distinct extents in two sexes, with many TEs silenced more in ovary and only a few silenced more in testis.

To explore the link between TE expression and piRNA programs in two sexes, we identified a set of 36 TE families repressed by piRNA pathway in at least one sex (see methods). For these TE families, there is a positive correlation between sex bias of piRNA production and sex bias of TE de-repression (Spearman’s *ρ*: 0.53, *P*<0.001; Figure 3A). For example, disruption of piRNA pathway by *rhi* mutations dramatically increases expression of three telomere-associated TEs (*HeT-A, TAHRE* and *TART*) in ovaries, where there are abundant piRNAs targeting these elements. On the contrary, much fewer piRNAs target these telomeric TEs in testes and expression of these TEs remained very low in *rhi* mutant males (Figure 3B; Figure 2A and 2E). This result indicates that telomeric TEs have a strong, intrinsic bias in their expression towards the female germline, and that piRNA pathway appears to have adapted to this bias generating respective antisense piRNAs in female, but not male, gonads. In contrast to ovary-biased TEs like telomeric elements, testis-biased TEs such as *Transib2* and *baggins1* are targeted by more antisense piRNAs in testis than ovary (Figure 3B; Figure 2A and 2E). Some TEs, such as *copia, mdg3* and *I-element* are strongly repressed by piRNAs in both sexes. For such elements, the sex bias in piRNA production does not always match that of TE repression (Figure 3A). Taken together, these findings suggest that, for most TEs, piRNA programs in males and females have adapted to differential TE activities between sexes.

**Figure 3.**
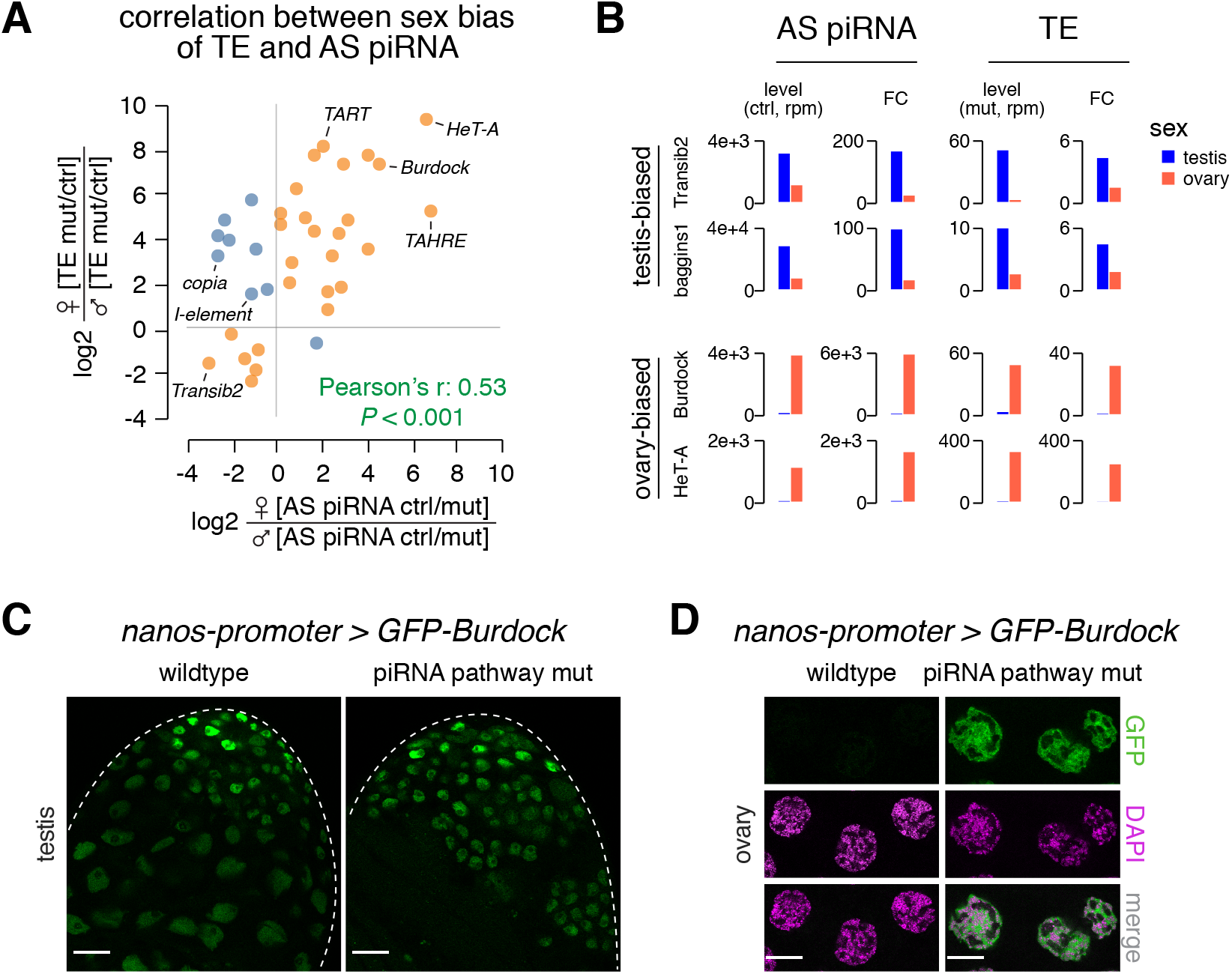
Sexually dimorphic piRNA programs parallel sex-specific TE expression. (A) Scatter plot displaying the correlation between sex biases of TE and TE-antisense piRNA. For each TE family, the loss of antisense piRNAs in *rhi* mutants was calculated in each sex (ctrl RPM over mut RPM). The sex bias of piRNAs was defined as the log2 ratio of piRNA loss in female over male. Similarly, TE de-repression in *rhi* mutants was calculated in each sex (mut RPM over ctrl RPM), and the sex bias was defined as the log2 ratio of TE de-repression in female over male. TE families that show a correlation between the sex bias of antisense piRNA and that of TE derepression are colored as orange, with the rest as blue. (B) Histograms showing profiles of two sex-biased TEs for each sex. Antisense piRNA levels refer to those in control gonads, TE levels refer to those in piRNA pathway mutants (*rhi^-/-^*), and the foldchange is calculated as mutant over control for TEs and the reverse for antisense piRNAs. (C) Confocal images of the apical tip of testis (left) and stage 7-8 nurse cells in ovary (right) that express a *Burdock-fused* GFP reporter. The reporter is expressed by *nanos* promoter that drives germline expression in both sexes, thus enabling the examination of piRNA silencing of *Burdock* sequences independent of natural expression patterns of *Burdock* transposon. Scale bars: 20μm.

To further explore whether differential expression of piRNAs between sexes has functional consequences, we studied *Burdock*, an LTR retro-transposon targeted by 53 times more piRNAs in ovary (3,756 RPM) than testis (70 RPM) (Figure 2A). We used a reporter composed of a fragment of *Burdock* expressed under the control of heterologous *nanos* promoter that drives expression in germline of both sexes (Handler et al., 2013). While reporter was efficiently silenced in ovaries of wild-type flies, it was strongly de-repressed in piRNA pathway mutants (*rhi^-/-^*) (Figure 3D), indicating that the piRNA program efficiently silences *Burdock* in female germline. In contrast, we observed strong reporter expression in testes of wild-type males, and the disruption of piRNA pathway in *rhi* mutants did not lead to an observable increase in its expression (Figure 3C). This finding shows that *Burdock* is not silenced in testes, likely as a result of very few *Burdock*-targeting piRNAs in males (Figure 2A). Notably, expression of endogenous *Burdock* is high in ovary (when piRNA production is disrupted) but low in both wild-type and mutant testis (Figure 2E; Figure 3B). Thus, similar to telomeric TEs, the ability of piRNA pathway to repress *Burdock* in female but not male germline correlates with an intrinsic bias for its expression in females. We conclude that differential expression of TE-targeting piRNAs in male and female gonads can have functional consequences in their abilities to silence TEs, suggesting a sexually dimorphic TE-silencing piRNA program operating in the gonad.

### Definition of piRNA clusters in testis with a new algorithm

To get deeper understandings of the piRNA program in male gonads, we sought to define the genomic origin of piRNAs and compare it between two sexes. Since genome-wide identification of piRNA clusters has only been done in ovary, we decided to systematically search for genomic loci that generate piRNA in testis. We noticed that two major clusters in testis identified to date, *Su(Ste)* and *AT-chX*, both contain internal tandem repeats, i.e., they are made of many copies of almost identical sequences (Aravin et al., 2001; Kotov et al., 2019). As a result, most piRNAs produced by these two loci mapped to the genome at multiple positions. However, the algorithm employed in previous studies to systematically define piRNA clusters in ovary only uses piRNAs that map to the genome at single unique positions (Brennecke et al., 2007; Mohn et al., 2014), raising the question of whether it is an appropriate approach to detect clusters like *Su(Ste)* composed primarily of internal tandem repeats. In fact, both *Su(Ste)* and *AT-chX* clusters were initially identified by different approaches (Aravin et al., 2001; Nishida et al., 2007).

Even though piRNAs produced from *Su(Ste)* and *AT-chX* cannot be mapped to single unique genomic loci, most of them mapped to several local repeats inside the respective clusters but nowhere else in the genome (Figure 4A). Taking advantage of this property, we developed a new algorithm that takes into account local repeats to define piRNA clusters (Figure S4A and S4B). Briefly, in addition to uniquely-mapped piRNAs, the algorithm searches for piRNA sequences that map to multiple positions within a single genomic region but nowhere else in the genome. This approach ensures that the identified region as a whole generates piRNAs, though the exact origin within the region remains unknown. Unlike the previous approach that uses exclusively uniquely-mapped piRNAs, this algorithm successfully identified *Su(Ste)* and *AT-chX*, two major piRNA clusters in testis that contain local repeats (Figure 4E and 4F).

**Figure 4.**
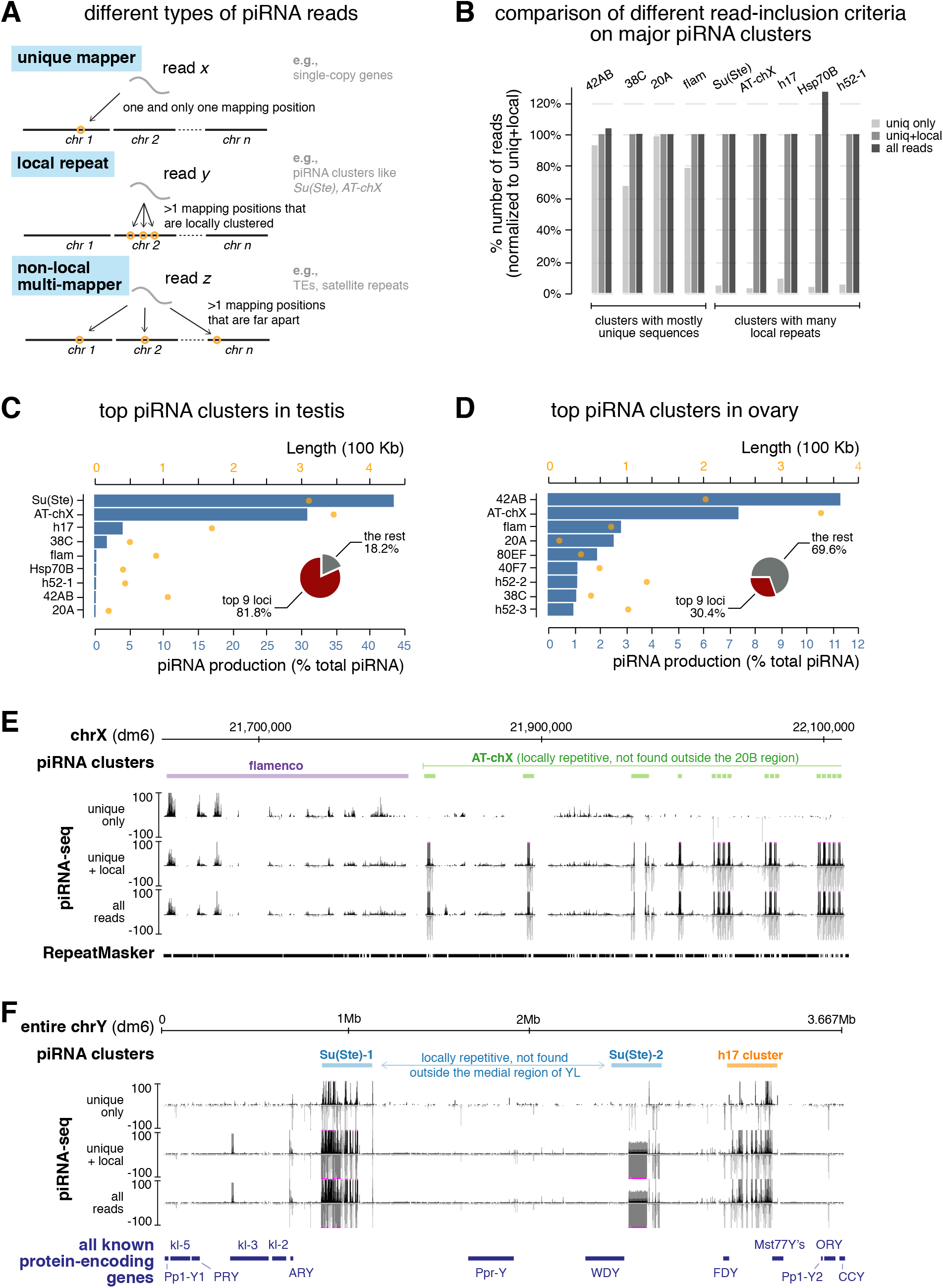
Definition of piRNA clusters in testis and ovary using a new algorithm. (A) Three types of piRNA reads, defined based on their mapping positions. Uniquely-mapped reads can be mapped to only one position in the genome and their origin is unambiguous. Reads derived from local repeats can be mapped to several positions in the genome; however, all of these mapping positions are locally clustered in a single genomic region. On the other hand, nonlocal multi-mappers can be mapped to multiple positions that are not restricted to one genomic region (typically mapped to more than one chromosome). Previously, only uniquely-mapped reads were used to define piRNA clusters and quantify their expression, as the genomic origin of multi-mappers is ambiguous. Inclusion of multi-mappers derived from local repeats, as shown in this study, allows identification of new piRNA clusters as well as a more accurate quantification of piRNA production from known clusters. At the same time, it preserves the certainty that reads are generated from genomic loci in question. See Figure S4 for detailed pipeline. (B) Histogram comparing numbers of mapped reads for major piRNA clusters using different readinclusion criteria as defined in (A). For each cluster, the number of mapped reads generated by different methods is normalized to the method that includes both unique and local repeat reads (the middle column). See also Figure S4 and methods. (C) Expression of the top 9 most active piRNA clusters in testis. Blue bars depict the contribution of each cluster to total piRNAs (%) and orange dots show cluster lengths according to dm6 genome assembly. Insert is a pie chart of the contribution of top 9 loci to total piRNAs in testis. (D) Same as in (C) but for ovary. (E) UCSC genome browser view of a peri-centromeric region (chrX) encompassing the entire *flamenco* locus (purple) and the distal part of *AT-chX* piRNA cluster (green). Below genomic coordinates (dm6) are piRNA coverage tracks using different read-inclusion criteria as defined in (A). Note that, whereas *flamenco* produce piRNAs that can be mostly mapped to unique genomic positions, *AT-chX* generates piRNAs that map to local repeats in this cluster, but nowhere else in the genome. Addition of non-local multi-mapper reads does not change the profile, indicating that the vast majority of piRNAs produced by the cluster are captured by unique+local mappings. (F) UCSC genome browser view of the entire Y chromosome that harbors two *Su(Ste*) loci (blue) and the novel *h17* piRNA cluster (orange). piRNA coverage tracks using different read-inclusion criteria are shown below genomic coordinates (dm6). At the bottom, all known Y-linked proteinencoding genes are drawn for reference (not to exact scale). Note that piRNA profiles of *Su(Ste*) and *h17* clusters collapse if piRNAs derived from local repeats are excluded.

We applied this new algorithm to systematically identify piRNA clusters active in testes. We recovered piRNA clusters known to be active in testes as well as piRNA clusters previously defined in ovaries (e.g., *42AB, 38C, 20A* and *flam*) (Figure 4C; Table S1). Furthermore, our search identified several novel piRNA loci. One of the novel piRNA clusters is located on Y chromosome flanked by *FDY* and *Mst77Y* genes (Figure 4C and 4F), which we called *h17* cluster using heterochromatin banding numbers (Gatti and Pimpinelli, 1983). Another novel locus is *h52-1*, flanked by *eIF4B* and *CG17514* genes on chr3L. *h52-1* harbors tandem local repeats composed of nested TE fragments that cannot be found elsewhere in the genome. Similar to piRNA clusters identified in ovaries, only a few clusters active in testes produce piRNAs from one genomic strand (e.g., *flam* and *20A*, so-called ‘uni-strand clusters’), and the majority are dual-strand clusters that generate piRNAs from both genomic strands (Figure 4E). In sum, our algorithm successfully found previously known piRNAs clusters and identified novel ones in *Drosophila* testes.

To compare new algorithm with the approach that considers only uniquely-mapped piRNAs, we applied both techniques to analyze the same testis piRNA dataset. This comparison showed that major piRNA clusters in testis can be divided into two groups (Figure 4B). The first group (*42AB, 38C, 20A* and *flam*) contains piRNA clusters that harbor mostly unique sequences, so including local repeats or even all reads do not substantially change their identification and quantification. On the other hand, the second group of genomic loci (*Su(Ste), AT-chX, h17, Hsp70B* and *h52-1*) is composed of piRNA clusters that contain many local repeats, and, accordingly, our new algorithm identified more than 10-fold more piRNAs produced from these loci (Figure 4B). Importantly, mapping of all piRNA reads (i.e., including multi-mappers not captured by our algorithm) only adds a negligible amount of piRNAs to these clusters, except for *Hsp70B* (Figure 4B). Thus, this algorithm is not only useful for finding new piRNA source loci but also provides a more accurate quantification of piRNA production from previously known clusters.

### Sex difference in piRNA cluster expression

To compare the expression of piRNA clusters between sexes, we first applied our algorithm to published ovary piRNA datasets (Figure 4D; Table S1) (ElMaghraby et al., 2019). Thus, piRNA clusters were defined and their activities were quantified in both sexes using the same algorithm, allowing for fair comparison. Surprisingly, our analysis revealed that *AT-chX*, originally described as a piRNA cluster in testes, is also highly active in ovaries. *AT-chX* locus consists of local repeats (Kotov et al., 2019), so piRNAs produced from this locus were excluded in previous studies that analyzed only uniquely-mapped reads. In fact, *AT-chX* is the second most active piRNA cluster in ovary, producing ~7% of total piRNAs.

Comparison between piRNA clusters in males and females revealed a clear sex difference: a small number of loci produce the majority of piRNAs in testis, which is not the case for ovary (Figure 5A). The two most active piRNA clusters in testes, *Su(Ste)* on Y chromosome and *AT-chX* on X chromosome, produce ~43% and ~31% of total piRNAs in testes, respectively (Figure 4C). They are followed by the novel piRNA cluster on Y chromosome, *h17*, that produces ~4% piRNAs. Along with another 6 loci, the top 9 piRNA clusters in testis account for 81.8% of total piRNAs. In comparison, only 30.4% of total piRNAs are made from the top 9 clusters in ovary, with the most active locus *42AB* producing ~11% of total piRNAs (Figure 4D). Whereas a few loci dominate the global piRNA population in testis, the ovary piRNA profile is shaped by many loci producing piRNAs in comparable amounts.

**Figure 5.**
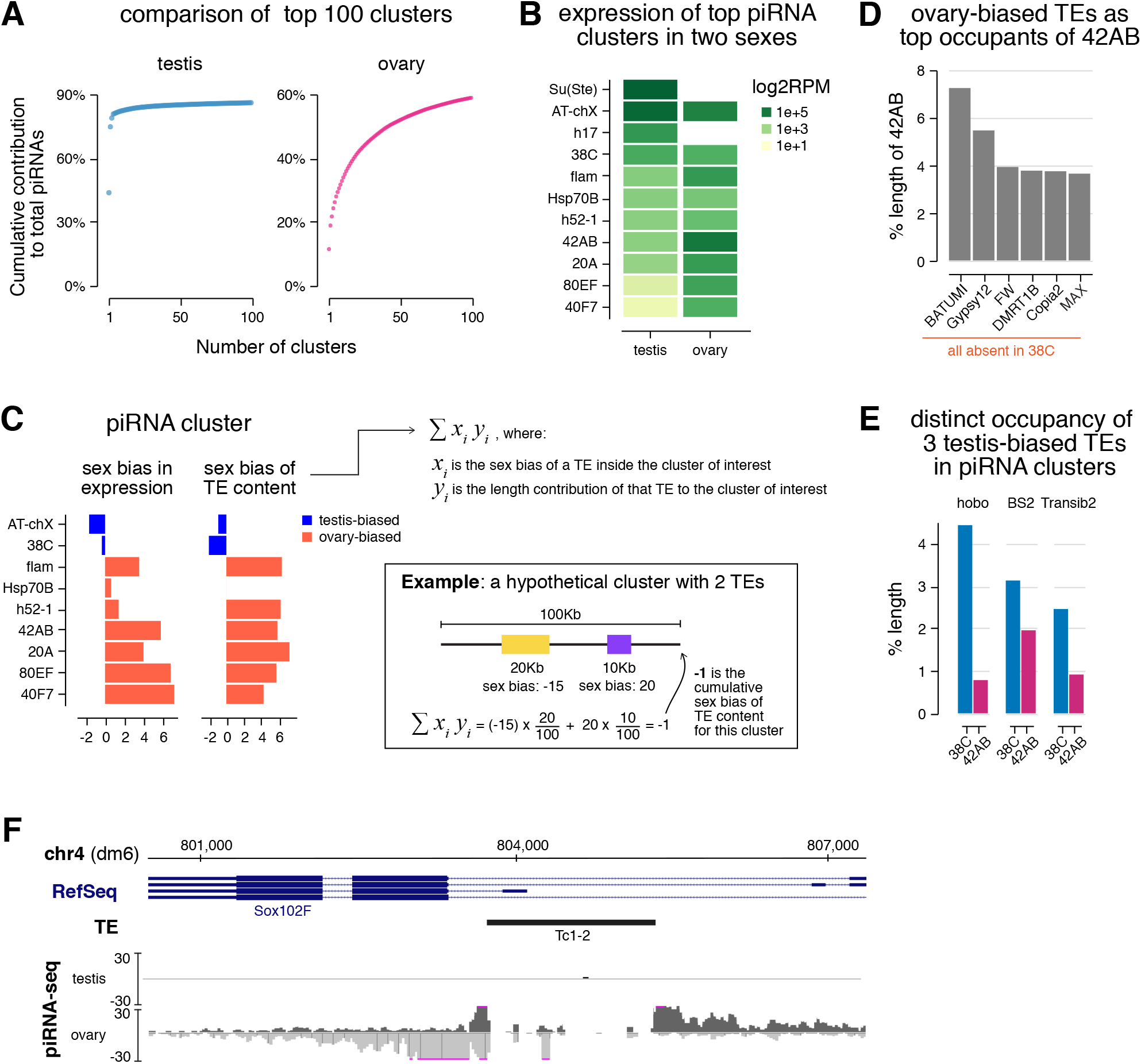
piRNA clusters are differentially employed to tame sex-specific TE expression. (A) Plot showing the cumulative contribution of top piRNA clusters to the total piRNA populations in testis (left) and ovary (right), up to 100 clusters. (B) Heatmaps showing piRNA production from major piRNA clusters. Note that *Su(Ste*) and *h17* clusters are Y-linked so there are no piRNAs from these loci in females that lack Y chromosome. (C) Bar graphs displaying the sex bias of piRNA cluster expression (left) and cumulative sex bias of the TE context for each cluster (right). Sex bias of piRNA cluster expression is defined as the log2 ratio of piRNA cluster expression in ovary over testis shown in (B), so ovary-biased ones are positive in value. Cumulative sex bias of cluster TE content is calculated by summing the sex bias of TEs (as described for Figure 2E) weighted by their length contributions to the cluster (equation shown on the right). An example is shown on the bottom right for a hypothetical cluster composed of two TEs with lengths and sex biases labeled accordingly for illustration. Only TEs showing strong sex biases were used in calculation. See also methods. (D) TE composition of ovary-biased *42AB* cluster. Shown are fractions of *42AB* cluster occupied by sequences from top 6 TE families. These 6 TEs are completely absent in *38C*, a testis-biased piRNA cluster. Expression of these 6 TEs is all ovary-biased (Figure S3A). (E) Contributions of 3 testis-biased TEs (Figure S3A) to the ovary-biased *42AB* cluster and testis-biased *38C* cluster. These TEs were selected as the most enriched by length in *38C* compared to *42AB*. (F) The *Sox102F* gene generates piRNAs in ovary, but not in testis. This locus harbors a single autonomous TE, *Tc1-2*, that has ovary-biased expression (Figure S3A). piRNA coverage tracks show both uniquely-mapped and local repeat-derived reads.

Next, we compared expression levels of different piRNA clusters in male and female gonads. Females lack Y chromosome, so they do not have piRNAs produced by Y-linked *Su(Ste)* and *h17* clusters. For major clusters present in both male and female genomes, we observed pronounced sex differences (Spearman’s *ρ* = 0.07; Figure 5B). For instance, *38C* produces more piRNAs than *42AB, 80EF* and *40F7* in testes, but the opposite trend is found in ovaries. Some loci such as *Sox102F* on chr4 (Mohn et al., 2014; Zhang et al., 2014) appear to be active only in ovaries but not in testes (Figure 5F). These differentially expressed piRNA clusters located on autosomes, which both males and females have two copies, exemplify the sex-specific usage of piRNA loci. Moreover, we examined expression levels of major piRNA clusters on chrX (*AT-chX, flam* and *20A*), which females have two copies (XX) and males have only one (XY). We found that a larger fraction of piRNAs originate from *AT-chX* in testes than ovaries, but the reverse was found for *flam* and *20A*, suggesting that copy numbers of piRNA clusters do not correlate well with their expression. Altogether, these findings illustrate a sexually dimorphic employment of piRNA clusters, where different loci are engaged differentially in a sex-specific manner.

Different piRNA clusters have distinct TE contents, so their differential expression might sculpt sex-specific piRNA programs with distinct TE-silencing capacities in males and females. To explore a link between the expression of piRNA cluster and its TE content, we computed cumulative sex bias of the TE content of each major piRNA cluster (Figure 5C). This was done by summing sex biases of individual TEs in the piRNA cluster weighted by their length contributions to the cluster (see example in Figure 5C). The sex bias of cluster TE content matches the sex bias in piRNA cluster expression, suggesting a link between the expression of piRNA clusters and TEs they control. To substantiate this finding, we analyzed sequence compositions of three differentially expressed piRNA clusters: *42AB* (ovary-biased), *38C* (testis-biased) and *Sox102F* (ovary-specific). The top 6 TEs most enriched by length in ovary-biased *42AB (batumi, gypsy12, FW, DMRT1b, copia2* and *max*) are all ovary-biased in their expression (Figure 5D; Figure S3A). Importantly, these 6 TEs are completely absent in testis-biased *38C* cluster. In contrast, three testis-biased TE families, *hobo, BS2* and *Transib2*, are more enriched in *38C* than in *42AB* (Figure 5E; Figure S3A). Moreover, ovary-specific *Sox102F* cluster harbors a single autonomous transposon, *Tc1-2*, which has higher activity in ovary (Figure 5F; Figure S3A). These examples show that differential expression of piRNA clusters in two sexes often matches the differential activities of TEs they control, supporting the notion that piRNA clusters are employed in a sex-specific fashion to cope with distinct TE landscape in male and female gonads.

### piRNA clusters composed of local repeats produce piRNAs that target host genes

Our analysis indicated that 13.8% of testis piRNAs might potentially be involved in targeting host genes as they can be mapped to protein-encoding genes in antisense orientation with a small number (0 to 3) of mismatches between piRNA and gene sequences (Figure 1B). To understand the genomic origin of these piRNAs, we further analyzed sequence compositions of piRNA clusters. We found that two clusters, *Hsp70B* and *h17*, both of which contain local repeats, generate piRNAs that have the potential to target host genes.

The *Hsp70B* cluster spans ~35Kb between two paralogous *Hsp70B* genes on chr3R, and it is active in both ovary and testis (Figure 6A). The body of *Hsp70B* cluster contains several TEs. Even though there are piRNAs mapping to these TEs, they can be mapped elsewhere in the genome as well, rendering it impossible to be certain that they originate from *Hsp70B* locus. In fact, this cluster was previously identified through the presence of uniquely-mapped piRNAs from flanking non-repetitive genes (Mohn et al., 2014). However, our algorithm that takes into account local repeats revealed piRNAs generated from a ~354bp local repeat at *Hsp70B* locus, which occupies nearly all inter-transposon space within this cluster. Importantly, these piRNAs mapped exclusively to this local repeat at *Hsp70B* cluster but nowhere else in the genome. Intriguingly, these repeats have a ~92% sequence identity to an exon of the *nod* gene, which encodes a kinesin-like protein necessary for chromosome segregation during meiosis (Carpenter, 1973; Hawley and Theurkauf, 1993; Zhang et al., 1990). *Hsp70B* cluster generates piRNAs that are antisense to *nod* with a 91.3% averaged nucleotide identity to it. This level of sequence similarity is close to that between *Suppressor of Stellate* piRNAs and their *Stellate* targets, the first known case of piRNA repression (Aravin et al., 2001; Vagin et al., 2006), suggesting that piRNAs produced from *Hsp70B* locus might be able to repress the *nod* gene.

**Figure 6.**
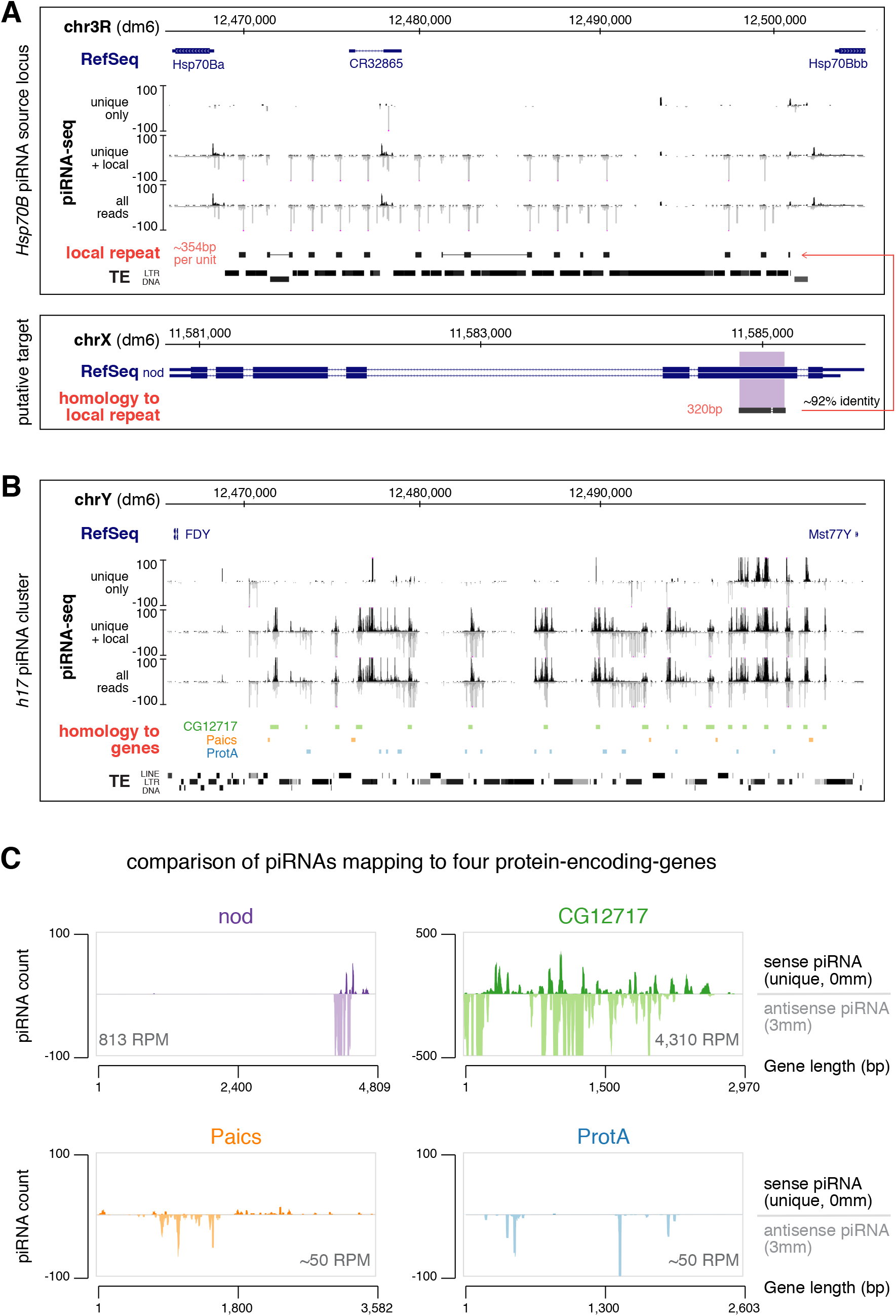
*Hsp70B* and *h17* piRNA clusters encode piRNAs that target host genes. (A) *Hsp70B* piRNA cluster (top) and the putative target, *nod* (bottom). piRNA coverage tracks using different read-inclusion criteria are shown below RefSeq and genomic coordinates (dm6) for *Hsp70B* cluster. ~354bp local repeats homologous to a 320bp exonic region of *nod* are depicted as solid blocks, which fill up most inter-TE space at this locus. Note that “unique+local” piRNA track does not include TE-derived piRNAs that map outside this locus, but it picks up *bona fide* local repeats that are homologous, but not identical, to *nod*. (B) *h17* piRNA cluster on Y chromosome. piRNA coverage tracks using different read-inclusion criteria are shown. Sequences with high levels of sequence similarity to protein-encoding genes are depicted as colored blocks (not to exact scale): *CG12717* (green), *Paics* (orange), *ProtA* (blue). Note that gene-homologous islands fill up most inter-TE space at this locus. Genomic coordinates are based on dm6 genome assembly. (C) Coverage of sense (unique, 0 mismatch) and antisense piRNAs (with up to 3 mismatches) over four putative, protein-encoding gene targets of testis piRNAs. Antisense piRNA abundance is shown for each gene.

The second locus producing piRNAs that might target host genes is the novel piRNA cluster *h17* on Y chromosome, which is only present in XY males (Figure 6B). This cluster spans more than 200Kb and includes two loci duplicated from chr2L and chrX, respectively, that contain almost the entire *CG12717* gene (which encodes a SUMO protease) and small parts of *Paics* (which encodes an enzyme involved in purine biogenesis) and *ProtA* (which encodes protamine, a sperm chromatin protein) (Mendez-Lago et al., 2011). These gene-homologous sequences are further duplicated locally on Y to over 20 copies and take up nearly all space in between TEs at *h17* cluster (Figure 6B; Figure S5B). However, these gene-related sequences likely do not retain coding potentials as they are frequently interrupted by TE sequences. *h17* cluster produces piRNAs antisense to *CG12717, Paics and ProtA* genes, with averaged levels of nucleotide identity 92.5%, 93.9%, and 91.0%, respectively. Together, two piRNA clusters, *Hsp70B* and *h17*, encode piRNAs with the potential to target both TEs and host genes.

We quantified expression of piRNAs antisense to *nod, CG12717, Paics* and *ProtA* genes from these two clusters. Even though these piRNAs all possess over 90% identity to their putative targets, their abundances differ dramatically (Figure 6C). *CG12717* gene is targeted by abundant piRNAs (4,310 RPM), comparable to the 15^th^ most targeted TE family in testis. piRNAs against *nod* are expressed at 813 RPM (~5-fold less compared to *CG12717*), while the levels of piRNA against *Paics* or *ProtA* are low (both ~50 RPM). In addition, nearly the entire length of *CG12717* gene is targeted by piRNAs, whereas only small parts of *nod, Paics* and *ProtA* are targeted. These findings suggest that *CG12717* and *nod* might be regulated by piRNAs in testis.

### piRNA-guided repression of SUMO protease *CG12717/pita* during spermatogenesis

To examine the role of piRNAs in gene regulation, we employed RNA-seq to analyze expression of host genes in testes of three different piRNA pathway mutants: *aub, zuc* and *spn-E* (Nishida et al., 2007; Pane et al., 2007; Schmidt et al., 1999; Stapleton et al., 2001). Transcriptome profiling revealed that only two genes, *CG12717* and *frtz*, exhibited ≥2-fold up-regulation in all three piRNA pathway mutants (Figure 7A). Unlike *CG12717*, there are very few, if any, antisense piRNAs targeting *frtz*, so its up-regulation likely reflects a secondary phenotype following TE de-repression. Strikingly, expression of *CG12717* increased more than 10-fold in all three mutants (Figure 7C), indicating that it is indeed strongly repressed by the piRNA pathway. Meanwhile, we observed no statistically significant up-regulation of *nod, Paics* or *ProtA* in these three mutants (Figure 7C), correlating with fewer piRNAs against these genes than *CG12717* (Figure 6C). While *CG12717* is expressed at a very low level in testes of wild-type males, it is highly expressed in ovaries (Figure 7D), consistent with the fact that *CG12717*-targeting piRNAs are encoded on Y chromosome. Thus, transcriptome profiling identifies *CG12717* as a target of piRNA repression and suggests that abundant antisense piRNAs with high target coverage might be required for efficient silencing.

**Figure 7.**
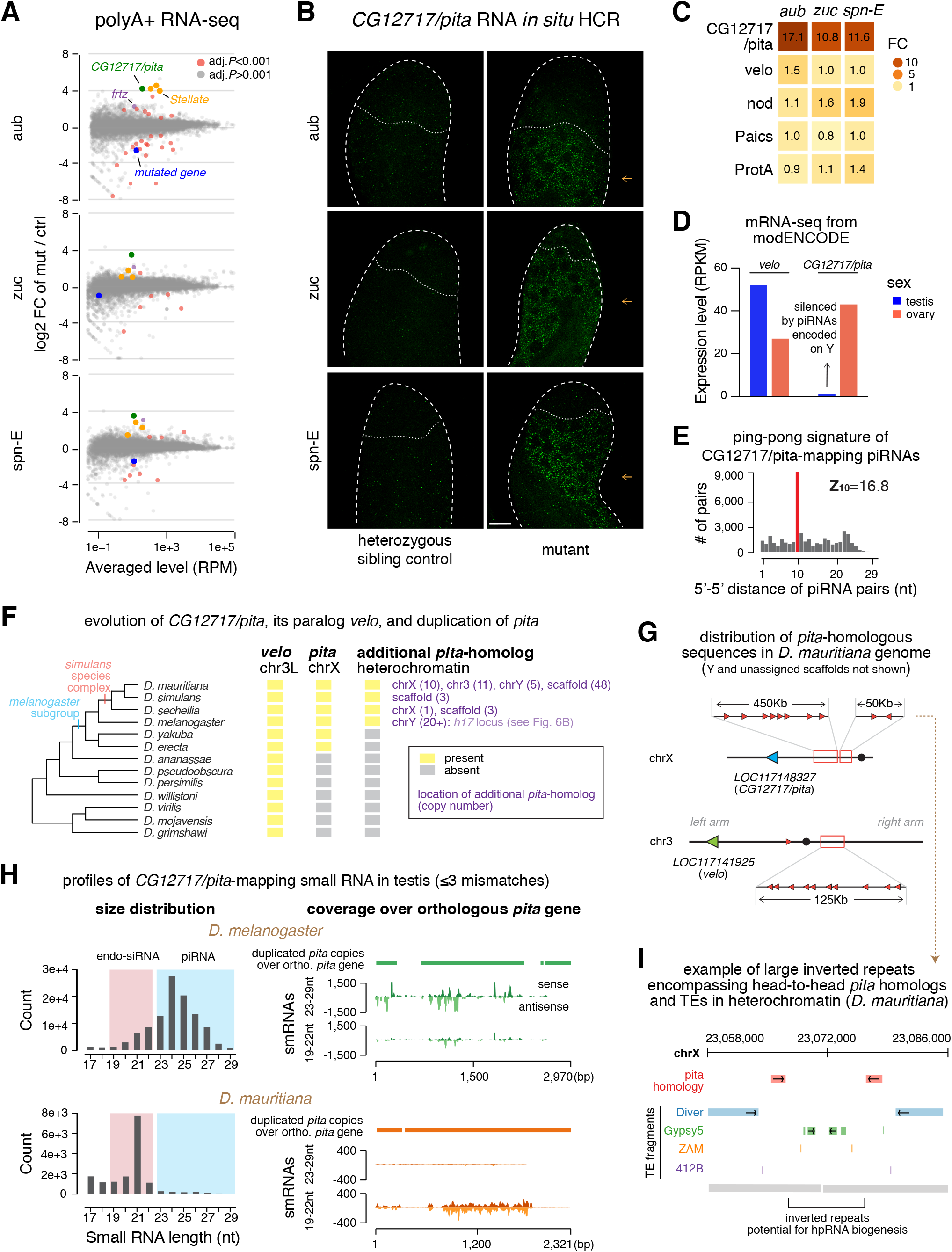
Regulation of *CG12717/pita* by small RNA pathways. (A) MA plots showing gene expression changes from polyA+ RNA-seq of *aub* (top), *zuc* (middle) and *spn-E* (bottom) mutant testes versus heterozygous sibling controls. Genes are marked red when passing a stringent statistical cutoff (adjusted *P*<0.001, from DESeq2). Additional coloring includes: *CG12717/pita* (green), annotated *Stellate* transcripts (orange), *frtz* (purple), and the mutated gene in each mutant (blue). (B) Confocal images of *pita* mRNAs detected by *in situ* HCR in *aub* (top), *zuc* (middle) and *spn-E* (bottom) mutant testes along with respective heterozygous sibling controls. Probes were designed against a ~400bp sequence unique to *pita* and absent on Y (Figure S5B), so they do not target *h17* piRNA precursors. Note that de-repression of *pita* in piRNA pathway mutants is observed specifically in differentiating spermatocytes (pointed to by orange arrows). Scale bar: 20μm. (C) Heatmaps showing fold-change of five protein-encoding genes in three mutant testes according to polyA+ RNA-seq shown in (A). (D) Bar graphs displaying modENCODE data of *pita* and its paralog *velo* expression in *D. melanogaster* gonads of both sexes. (E) Analysis of ping-pong processing of *pita*-mapping piRNAs. Histogram shows distribution of 5’-to-5’ distances of complementary piRNA pairs with an enrichment for 10nt (i.e., ping-pong signature). To select secondary piRNAs processed from *pita* transcripts, only reads that map perfectly to *pita* mRNAs in sense orientation and do not map perfectly to *h17* cluster were used in this analysis. Antisense piRNAs were selected allowing up to 3 mismatches. (F) Cladogram of major species in *Drosophila* genus (left) and the evolutionary history of *velo, pita* and *pita*-related sequences in genomes of these species (right). Orthologs were identified based on sequence homology and synteny. Shown in purple are locations of additional *pita* copies in each species and copy numbers in parenthesis. (G) Cartoon depicting distribution of *pita*-homologous sequences in *D. mauritiana* genome. Orthologous *pita* is marked blue, orthologous *velo* is marked green, and the duplicated, candidate sources of *pita*-targeting endo-siRNAs are marked red. Note that they scatter across peri-centromeric heterochromatin of chrX and chr3, as well as chrY and scaffolds (not shown). (H) Profiles of *pita*-mapping small RNAs in testes of *D. melanogaster* (top) and *D. mauritiana* (bottom). Size distributions are shown on the left. Coverage plots over the orthologous *pita* gene in each species are shown on the right, including: cumulative alignment of heterochromatic, duplicated copies of *pita* over the syntenic, orthologous *pita* (top, solid bar), stranded coverage of 23-29nt piRNAs (middle, histogram) and 19-22nt endo-siRNAs (bottom, histogram) over the orthologous *pita* gene. (I) Illustration showing two representative head-to-head copies of *pita* homology (red) in the peri-centromeric heterochromatin of *D. mauritiana* X chromosome. *pita*-related sequences are flanked by TEs and are part of a large inverted repeat that could potentially permit hpRNA biogenesis.

piRNA-guided cleavage of target RNAs often triggers the production of secondary piRNAs from target RNAs in a process dubbed ping-pong cycle (Brennecke et al., 2007). Examination of piRNA sequences revealed abundant piRNAs derived from the entire length of *CG12717* mRNAs (Figure 6C). In contrast, we found few piRNAs processed from transcripts of *nod, Paics* or *ProtA*. Furthermore, sense piRNAs derived from *CG12717* mRNAs and antisense piRNAs produced from *h17* cluster demonstrated a strong ping-pong signature (Z_10_=16.8; Figure 7E), characteristic of active ping-pong cycle. This finding further suggests the direct cleavage of *CG12717* transcripts guided by *h17* piRNAs. Finally, we performed RNA *in situ* hybridization chain reaction (*in situ* HCR) to examine *CG12717* expression. Expression of *CG12717* is very low in wild-type testis, but it was significantly increased in testes of *aub, zuc* and *spn-E* mutants (Figure 7B). Upon release of piRNA silencing in these three mutants, *CG12717* is specifically expressed in differentiating spermatocytes, but not in germline stem cells or mitotic spermatogonia. Interestingly, *Stellate* is expressed at the same stage when the silencing by *Su(Ste*) piRNAs is removed (Aravin et al., 2004). As our results indicate that expression of *CG12717*, a SUMO protease gene related to Ulp2 in yeast (Berdnik et al., 2012), is strongly repressed by piRNAs, we propose to name it *pita* (<u>pi</u>RNA <u>ta</u>rget).

To understand how piRNA-dependent regulation of *pita* has evolved, we performed a blastN search using *Drosophila melanogaster pita* gene against genomes of other *Drosophila* species. We found multiple copies of *pita*-related sequences in genomes of *Drosophila simulans* species complex (*D. simulans, D. sechellia* and *D. mauritiana*) (Figure 7F), but not in more distantly related species like *D. erecta* or *D. yakuba*. Similar to *h17* cluster in *D. melanogaster*, these *pita*-related sequences reside in TE-rich regions (either peri-centromeric heterochromatin or unassigned scaffolds) in *D. simulans* species complex. While all *pita*-related sequences are exclusively located at *h17* on Y chromosome of *D. melanogaster, pita*-homologous sequences can be found on different chromosomes in genomes of *D. simulans* species complex. For instance, in *D. mauritiana, pita*-homologous sequences can be found on at least chrY, chrX, chr3L and chr3R (Figure 7G). Therefore, duplications of *pita*-related sequences into heterochromatin have occurred in all four species.

To investigate whether heterochromatic *pita*-homologous sequences produce small RNAs in testes of other species, we analyzed published small RNA datasets from testes of *D. simulans* and *D. mauritiana* (Kotov et al., 2019; Lin et al., 2018). We found no small RNAs mapping to the orthologous *pita* gene in *D. simulans* testes, but abundant ones in *D. mauritiana* testes (Figure 7H). Unexpectedly, unlike 23-29nt *pita*-mapping piRNAs in *D. melanogaster, pita*-mapping small RNAs in *D. mauritiana* are mostly 21nt long, indicating that they are endo-siRNAs. These endo-siRNAs have on average 93.5% identity with the *D. mauritiana pita* gene. Notably, similar to other dual-strand piRNA clusters described in *D. melanogaster* ovaries (Czech et al., 2008; Le Thomas et al., 2014), *h17* cluster in *D. melanogaster* testes also generates *pita*-mapping endo-siRNAs, though much less abundant than 23-29nt piRNAs (Figure 7H). Examination of heterochromatic, *pita*-homologous sequences in *D. mauritiana* genome revealed that most of them are arranged head-to-tail (Figure 7G). However, there are four instances where *pita*-homologous sequences are arranged head-to-head (Figure 7G and 7I), which could potentially generate hairpin RNAs (hpRNAs), the preferred substrate for processing into endo-siRNAs by Dicer. Thus, targeting of *pita* by small RNAs in testis seems to be conserved in two *Drosophila* species. While *pita* is repressed mostly by piRNAs in *D. melanogaster*, it is targeted nearly exclusively by endo-siRNAs in *D. mauritiana*, suggesting two related but distinct regulation strategies employed in sibling species that diverged less than 5 million years ago.

In addition to *pita*, there is another SUMO protease, *veloren (velo*) in *D. melanogaster* genome. *pita* and *velo* are paralogs whose homologous domains share 75% nucleotide identity (Figure S5A). In agreement with the sequence similarity, functions of Pita and Velo in SUMO deconjugation pathway were shown to be partially redundant (Berdnik et al., 2012). Phylogenetic analysis showed that, while *velo* is found at syntenic locations on chromosome 3 throughout the *Drosophila* genus, *pita* is much younger and was only born after the split of *D. melanogaster* and *ananassae* species subgroups (Figure 7F). These results indicate that *pita* and *velo* have evolved from a common ancestor gene, via inter-chromosomal duplication from chr3 to X chromosome.

Considering the 75% nucleotide identity between the parts of *pita* and *velo* genes in *D. melanogaster, pita*-targeting piRNAs produced from *h17* cluster have a potential to target *velo* transcripts. However, we found that none of the *pita*-antisense piRNAs can be mapped to *velo* transcript perfectly. Moreover, ~200-fold fewer piRNAs have a potential to target *velo* with one to three mismatches. Transcriptome profiling in testes of *aub, zuc* and *spn-E* mutants showed that, unlike *pita, velo* is not repressed by piRNAs (Figure 7C). In addition, while *pita* is only expressed in ovaries, *velo* is expressed in both testes and ovaries and, in fact, has a higher expression level in testes (Figure 7D). These results show that Y-linked *h17* piRNAs repress specifically *pita*, but not its paralog, *velo*, suggesting that a high degree of complementarity is required for efficient piRNA silencing. Therefore, piRNAs distinguish closely related paralogs to achieve sex- and paralog-specific gene regulation.

Taken together, our results allowed us to reconstruct the evolutionary history of two paralogous, Ulp2-like SUMO protease genes. First, the *pita* gene was born via inter-chromosomal duplication after the split of *D. melanogaster* and *ananassae* species subgroups. This then permitted the differentiation of *velo* and *pita* functions, though these two genes remain in part functionally redundant in *D. melanogaster* (Berdnik et al., 2012). Next, divergence between *pita* and *velo* sequences created an opportunity for paralog-selective gene regulation by small RNA-guided mechanisms. This was achieved by duplications of *pita* sequences into heterochromatin in genomes of *D. melanogaster* and *D. simulans* species complex. It is plausible that, initially, heterochromatic, *pita*-homologous sequences did not play a role in gene regulation, as illustrated by the absence of *pita*-mapping small RNAs in *D. simulans*. However, subsequent expansion and interaction with TE sequences might have enabled the evolution of two distinct repression mechanisms, via production of *pita*-targeting piRNAs and endo-siRNAs, that dominated in *D. melanogaster* and *D. mauritiana*, respectively. Repression of *pita* by Y-linked piRNAs led to its specific repression in *D. melanogaster* testis, implicating the piRNA pathway in establishing distinct expression patterns of closely related paralogs after gene duplication.

## DISCUSSION

Previous studies systematically analyzed piRNA profiles in female gonads of *D. melanogaster*, revealing an essential role of piRNAs in regulation of many TEs (Brennecke et al., 2007; Li et al., 2009; Malone et al., 2009). However, these studies only provided a single snapshot of the relationship between TE and piRNA defense system, as they are insufficient to understand how the piRNA program might adapt to changing TE repertoire and different levels of their expression. To this end, several studies explored the piRNA pathway in other species of *Drosophila* (Malone et al., 2009; Rozhkov et al., 2010; Saint-Leandre et al., 2020). These studies revealed that piRNA profiles are different across species, suggesting an adaptation of the defense mechanism to distinct challenges. However, drastic differences in both TE contents and piRNA cluster sequences even among closely related *Drosophila* species (Kofler et al., 2015; Lerat et al., 2011; Malone et al., 2009) make it difficult to disentangle different factors that sculpt species-specific piRNA programs. Here, we examined TE expression in males and females of the same species, revealing strong differences in TE activities between sexes. This allowed us to compare piRNA programs in two sexes with similar genomic contents (except Y chromosome).

Another obstacle to understanding responses of the piRNA program to TEs is properly assessing TE expression. *D. melanogaster* genome includes over 100 different TE families whose expression levels can be measured by standard methods such as RNA-seq. However, TE expression in wild-type animals is greatly suppressed by the piRNA pathway (>100-fold for some families) (ElMaghraby et al., 2019). Therefore, in order to understand true expression potentials of TEs, it is necessary to study their expression upon removal of piRNA silencing, which is difficult to do in species other than model organisms like *D. melanogaster*. In this work, we examined the TE expression in piRNA pathway mutants, revealing genuine potentials of TE expression in both sexes. Combined analysis of TE and piRNA expression showed responses of the piRNA program to distinct TE expression profiles in two sexes.

Analysis of the genomic origin of piRNAs represents an important but challenging task. As piRNA sequences are short (23-29nt) and often derive from repetitive genomic regions, a large fraction of sequenced piRNA reads can be mapped to multiple genomic loci, preventing an unambiguous assignment of their origin. Accordingly, algorithms employed in previous studies only used the small fraction of piRNA reads that can be mapped to the genome at single unique positions to identify genomic regions that generate piRNAs. We took advantage of the fact that some genomic repeats are local, i.e., they reside within one genomic region and are absent in the rest of the genome, to develop a new algorithm for piRNA cluster definition and analysis (Figure 4A and Figure S4). This approach was successful in identifying new piRNA clusters. Furthermore, it also provided a more accurate quantification of the piRNA cluster expression. We found that *Hsp70B* cluster generates piRNAs against the *nod* gene. In addition, we discovered a novel cluster, *h17*, on Y chromosome that generates piRNAs against three host genes and ensures the strong silencing of SUMO protease, *CG12717/pita*, during spermatocyte differentiation.

Our identification of the novel *h17* cluster on Y expanded known functions of entirely heterochromatic Y chromosome (Figure 4F and Figure 6B). Three functionalities have been assigned to Y by the early 1980s (Gatti and Pimpinelli, 1983). First, together with X chromosome, Y encodes rDNA loci that express rRNAs and mediates meiotic pairing with X. Second, Y encodes six protein-encoding genes, so-called “fertility factors”, whose protein products are required for completion of spermatogenesis. Finally, Y chromosome harbors the *Su(Ste*) locus that generates piRNAs to suppress *Stellate* genes to safeguard normal spermatogenesis (Aravin et al., 2001; Vagin et al., 2006). A handful of new protein-encoding genes were discovered on Y in the past two decades (Bernardo Carvalho et al., 2009; Krsticevic et al., 2010), however, many of them appeared dispensable. Our finding that Y chromosome encodes a novel piRNA cluster and produces piRNAs to regulate expression of the *pita* gene assigns a new function to Y chromosome.

### Sexual dimorphism of TE expression and TE-silencing piRNA programs

*D. melanogaster* is an excellent model to study TE regulations and host-TE interactions, as its genome harbors many TE families that are transcriptionally and transpositionally active, generating new insertions in the population (Kofler et al., 2015). Our results indicate that expression of both TEs and piRNAs is sexually dimorphic. The majority of TE families are strongly expressed in ovaries, though some TEs are more active in testes. In line with this, our results indicate a stronger TE-silencing piRNA program in female gonads (Figure 2).

For TEs to be evolutionarily successful, they need to evolve strategies to maximize their chance to be inherited and expanded through generations. For example, TEs often hijack germline gene expression programs to be preferentially active in germ cells. Germline-biased expression leaves the choice of expression to either female or male germline, or both. Importantly, the two sexes employ distinct evolutionary strategies and have different contributions towards the zygote. While the major contribution of sperm is its genome, oocyte contributes large amounts of yolk, various protein factors, RNAs and organelles such as mitochondria, in addition to its genome. This sexual asymmetry in their contributions to the next generation has important implications for reproduction strategies of TEs. TEs active in the male germline need to complete the entire life cycle from transcription to genomic insertion before sperm maturation, in order to propagate. In contrast, once transcribed, TEs active during oogenesis could finish their life cycle in the zygote after fertilization, as long as transcribed TE transcripts are deposited into the oocyte. The latter strategy is also used by mammalian L1 retrotransposon that is expressed during gametogenesis, but genomic insertions might occur later during early embryogenesis (Kano et al., 2009). Thus, the expression bias towards ovaries observed for most TEs can be explained by an advantage for their proliferation, specifically, the extended window to finish their life cycle, in female germline.

There are a few TEs that bias testis for expression, suggesting that there are likely male-specific vulnerabilities exploitable by these elements. For example, male germ cells use a testis-specific gene expression machinery (e.g. tTAF and tMAC) to transcribe meiotic and post-meiotic genes (Beall et al., 2007; Hiller et al., 2004). TEs might exploit this tissue-specific transcriptional machinery to enable their sex-biased expression. It will be important in the future to uncover molecular mechanisms underlying differentially expressed TEs between sexes.

Analysis of piRNA profiles in testis and ovary indicates that piRNA programs have adapted to sex-biased TE expression (Figure 3). The most striking example is the nearly exclusive expression of telomeric TEs and corresponding antisense piRNAs in the female germline. Our results suggest that differential expression of piRNA clusters in two sexes together with differential TE-targeting capacity of each cluster contributes to the sex-specific, TE-targeting piRNA program. We found that piRNA cluster expression is sexually dimorphic. Besides the *Su(Ste)* locus, we identified another major cluster on Y chromosome that is only active in XY males. However, sex-biased expression is not restricted to Y-linked clusters, as many X-linked and autosomal clusters have differential activities between sexes as well. Besides differential expression, genomic analysis showed differences in piRNA cluster TE contents, suggesting that different piRNA clusters are, to some extent, specialized to target different sets of TEs. Importantly, sex bias in cluster expression and their TE-targeting potentials are linked: clusters preferentially targeting ovary-biased TEs are more active in ovary, while testis-biased clusters tend to target testis-biased TEs (Figure 5). Hence, piRNA clusters appear to be employed differentially by two sexes to counteract specific TE threats they face. What determines the differential expression of piRNA clusters between sexes awaits future studies. Previous work suggests that TE promoters embedded in piRNA clusters retain their activities (Mohn et al., 2014). Contribution of TE promoters to piRNA precursor transcription from piRNA clusters might explain the correlation between expression of clusters and their TE targets.

### Satellite DNA as target of piRNA silencing

Satellite DNAs can be classified as either simple or complex satellites based on the length of repeating units, and they occupy large portions of *Drosophila* genome, particularly at peri-centromeric and sub-telomeric regions (Hsieh and Brutlag, 1979; Karpen and Spradling, 1992; Larracuente and Presgraves, 2012; Lohe et al., 1993). We found piRNAs expressed from three major families of complex satellites: sub-telomeric *HETRP/TAS, Responder (Rsp)*, and *SAR/1.688* (including 359bp). In fact, piRNAs can be mapped to both strands of complex satellites in gonads of both sexes, and they often possess ping-pong signature (Figure 1C). Thus, our results expand the previous observation of piRNAs mapping to one strand of *Rsp* (Saito et al., 2006) and establish complex satellites as dual-strand piRNA clusters and potential targets of piRNA silencing in *Drosophila* germline of both sexes. Our analysis was focused on complex satellites, as simple satellite repeats are still largely intractable to sequencing technologies today (Khost et al., 2017). However, a recent study reported that transcripts from AAGAG simple satellite repeats regulate heterochromatin in male germline and are required for male fertility (Mills et al., 2019). It will be interesting to determine whether simple satellites produce piRNAs and, if so, whether their piRNA production is required for male fertility.

piRNAs loaded onto the nuclear Piwi protein guide heterochromatin assembly (Le Thomas et al., 2013; Rozhkov et al., 2013; Sienski et al., 2012; Wang and Elgin, 2011). For this reason, satellite piRNAs might play a role in establishing germline heterochromatin, similar to heterochromatin formation guided by siRNAs in fission yeast (Hall et al., 2002; Volpe et al., 2002). While the function of complex satellites remains mostly elusive, *Rsp* has been implicated in a meiotic drive system called segregation distortion (Hartl, 1973; Larracuente and Presgraves, 2012; Wu et al., 1988). During male meiosis, the *Segregation Distorter (SD*) allele enhances its own transmission to haploid cells at the cost of wild-type (*SD*^+^) allele in *SD/SD*^+^ heterozygous males, violating Mendelian law of inheritance. Importantly, segregation distortion requires the presence of a sufficient number of *Rsp* satellite repeats *in trans*. Though described more than 60 years ago (Sandler et al., 1959), the molecular mechanism of segregation distortion remains unknown. Intriguingly, mutations of *aubergine (aub)*, a PIWI protein, were found to be enhancers of *SD* (Gell and Reenan, 2013). Together with our data, these data suggest that piRNA pathway may play a role in segregation distortion during spermatogenesis.

### Regulation of host genes by piRNAs

Though the central and conserved function of piRNA pathway seems to be TE repression, other functions were also described in several organisms (reviewed in Ozata et al., 2019). The role of piRNAs in regulating host gene expression is particularly intriguing and remains somewhat controversial. The first described piRNAs, *Su(Ste)* piRNAs, silence the expression of *Stellate* genes (Aravin et al., 2001; Vagin et al., 2006). However, *Stellate* genes and their piRNA suppressors appear to resemble selfish toxin-antitoxin systems rather than representing an example of host gene regulation (Aravin, 2020). Since the discovery of piRNA pathway, there have been several studies reporting host protein-encoding genes regulated by *Drosophila* piRNAs (reviewed in Rojas-Ríos and Simonelig, 2018). In this work, we analyzed the ability of *Drosophila* piRNAs to regulate host genes in testes, by examining gene-targeting piRNAs and changes in host gene expression across three piRNA pathway mutants. We found piRNAs targeting four host genes: *nod* (a kinesin-like protein), *CG12717/pita* (a SUMO protease), *Paics* (a metabolic enzyme) and *ProtA* (a sperm chromatin protein). These four genes are targeted by antisense piRNAs generated from two piRNA clusters that contain sequence homology to them. However, only one of the four, *CG12717/pita*, is substantially repressed (over 10-fold) by piRNAs (Figure 7). As *pita*-silencing piRNAs are encoded on Y chromosome and thus only expressed in males, they are responsible for differential expression of *pita* in two sexes. Indeed, in wild-type files, *pita* is specifically silenced in male gonads while highly expressed in female counterparts. Thus, our results establish the ability of piRNAs to repress host protein-encoding genes, and, at the same time, suggest that this role is likely restricted to a small number of genes.

Our results indicate that piRNA-guided repression of host genes requires a sufficient number of targeting piRNAs. While all four genes are targeted by piRNAs with similar levels of sequence identity (91-94%, i.e., about 2 mismatches per piRNA), the abundance of piRNAs against each gene differs drastically. There are much more *pita*-targeting piRNAs than the other three gene targets, at a level comparable to the 15^th^ most targeted TE. Furthermore, while *pita* is targeted along almost the entire length, only small regions of other three genes are targeted by piRNAs. These differences in piRNA abundance and distribution of target sites could explain strong repression of *pita*, in contrast to the other three genes. It is possible that these genes are still regulated by piRNAs at specific stages, the question that remains to be further investigated. Importantly, abundant *pita*-silencing piRNAs do not repress the *pita* paralog, *velo*, that has a 75% sequence identity with *pita*, indicating that a high complementarity between piRNA and target may be important for efficient silencing. In agreement with these, a previous report indicated that a similar level of sequence identity (~76%) is insufficient for the silencing of *vasa* by *AT-chX* piRNAs (Kotov et al., 2019). Therefore, both high expression and high complementarity with targets might be required for efficient piRNA silencing in *D. melanogaster*.

This conclusion is important for analyzing the potential of piRNAs to repress host proteincoding genes. Unlike miRNAs, sequences of piRNAs are extremely diverse. Accordingly, if mismatches between piRNA and its target are well tolerated, a large number of cellular mRNAs should be targeted and repressed by piRNAs. Indeed, some host genes were proposed to be repressed by a few piRNA species that have multiple mismatches to mRNA sequences (Gonzalez et al., 2015; Klein et al., 2016; Rojas-Ríos et al., 2017; Saito et al., 2009). Our results suggest that such a spurious targeting by individual piRNAs is unlikely to cause repression. In fact, a high threshold for efficient target repression might permit production of diverse piRNA sequences against genuine targets such as TEs, without unintended interference with host gene expression.

### The role of piRNA in evolution

Analysis of *pita* repression revealed a remarkable picture of evolutionary innovation (Figure 7). piRNA-dependent repression of *pita* occurs in *D. melanogaster* but not in its sibling species, suggesting its rather recent origin. Efficient silencing of *pita* is linked to the presence of multiple copies of *pita*-homologous sequences in a piRNA cluster inside heterochromatin. Interestingly, duplications of *pita* sequences into, and their expansion within, heterochromatin can be found in three closely related species of *D. simulans* complex, in addition to *D. melanogaster*. However, distribution and copy number of *pita*-related sequences differ among these four species. In fact, both *h17* locus that generates *pita*-silencing piRNAs and its two flanking protein-encoding genes, *FDY* and *Mst77Y*, evolved after the split of *D. melanogaster* and *D. simulans* species complex (Carvalho et al., 2015; Krsticevic et al., 2010; Mendez-Lago et al., 2011), suggesting that the entire locus is unique to *D. melanogaster*. Furthermore, no small RNAs are generated from heterochromatic *pita* sequences in *D. simulans*, while endo-siRNAs are made against *pita* in *D. mauritiana*. The neutral theory of molecular evolution provides the most parsimonious interpretation of these results. This theory suggests that the initial duplication of *pita* sequences into heterochromatin might have been a random event that did not play a role in regulating the ancestral *pita* gene. However, subsequent evolution of *pita*-related sequences inside heterochromatin gave rise to two different modes of regulations, piRNA and endo-siRNA, in two different but closely related species. The emergence of small RNA-mediated repression was probably facilitated by the fact that *pita* itself was recently evolved and retains partially redundant functions with its paralog, *velo* (Berdnik et al., 2012), allowing independent regulation of two paralogs.

The evolutionarily innovative role of piRNAs in regulating host genes in *Drosophila* has interesting parallels in other organisms. Pachytene piRNAs expressed during spermatogenesis in mammals evolved very fast and are generally poorly conserved (Özata et al., 2020). The function of pachytene piRNAs is under active debate as no obvious targets can be easily discerned by analysis of their sequences (Aravin et al., 2006; Girard et al., 2006). Recently, knockout of one pachytene piRNA cluster led to unexpected conclusion that a small fraction of piRNAs promote biogenesis from other piRNA clusters and regulate the expression of a few host genes, while the vast majority do not target any transcripts (Wu et al., 2020). Thus, mammalian pachytene piRNAs can be considered a selfish system that occasionally involves in regulation of the host gene expression. Species-specific regulation of host genes by piRNAs in both *Drosophila* and mouse suggests that piRNA pathway is used in evolution to create innovation in gene regulatory networks that might contribute to speciation. More generally, piRNAs might promote the evolvability of animal species. Though it is difficult to establish the function of any molecular mechanism in evolution, this proposal makes a testable prediction that host genes repressed by piRNAs differ even among closely related species. Future studies in non-model organisms will shed light on the role of piRNAs in evolution and speciation.

## ACKNOWLEDGEMENT

We are grateful to William Theurkauf, Trudi Schüpbach, Julius Brennecke and Bloomington Drosophila Stock Center for fly stocks. We thank Katalin Fejes Toth and members of Aravin lab for discussion and comments. We appreciate the help of Maria Ninova and Fan Gao (Bioinformatics Resource Center, Caltech) with bioinformatics analysis, the help of Grace Shin and Maayan Schwarzkopf with HCR experiments, the help of Giada Spigolon and Andres Collazo (Biological Imaging Facility, Caltech) with microscopy, and the help of Igor Antoshechkin (Millard and Muriel Jacobs Genetics and Genomics Laboratory, Caltech) with sequencing. This work was supported by grants from the National Institutes of Health (R01 GM097363) and by the HHMI Faculty Scholar Award to A.A.A.

## SUPPLEMENTARY ITEM TITLES AND LEGENDS

**Figure S1.**
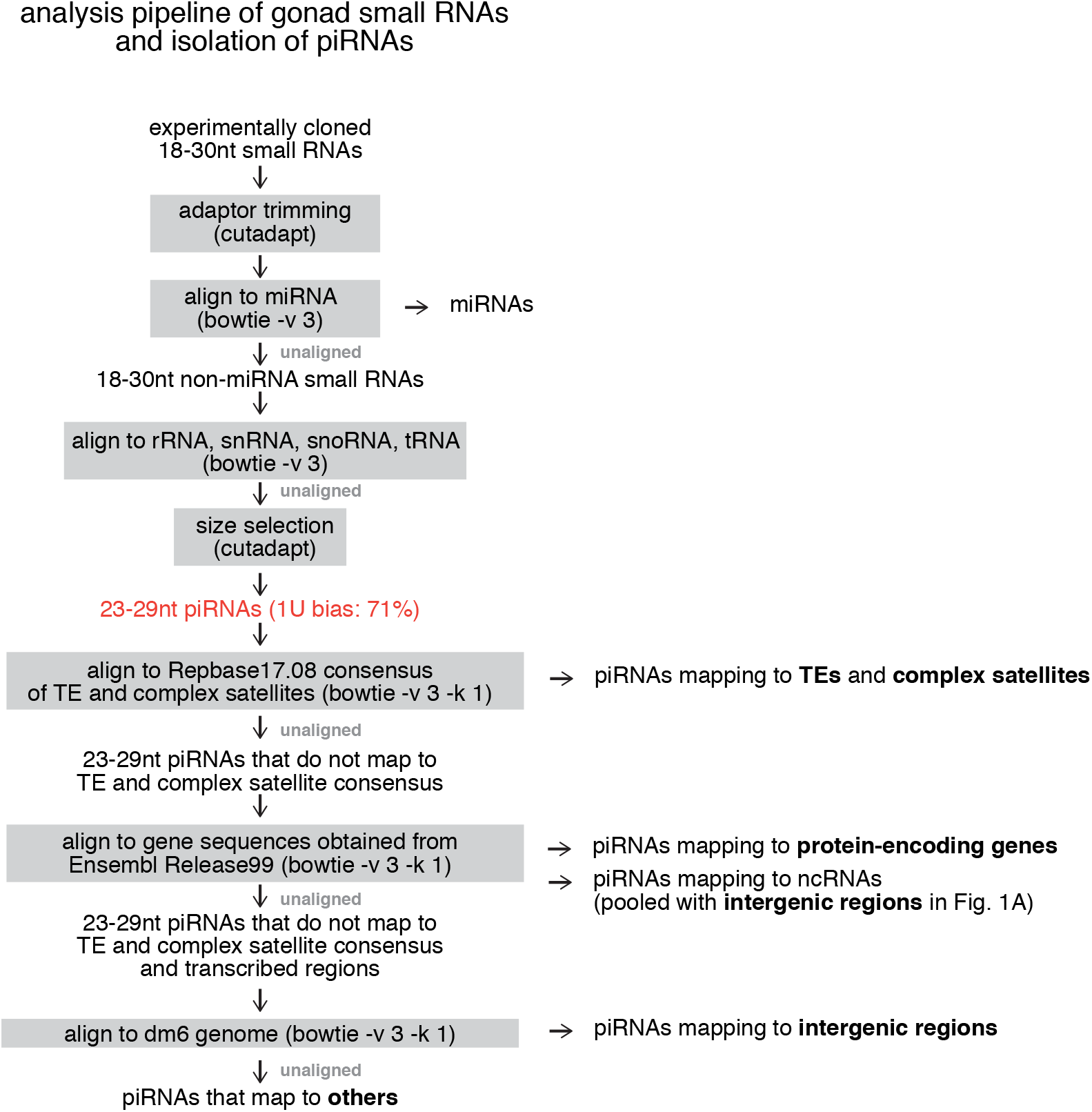
Analysis pipeline of gonad small RNAs. Related to Figure 1. Flow chart showing step-wise isolation of piRNAs from total small RNAs and subsequent mappings to different annotations (repeats, protein-encoding genes and genome).

**Figure S2.**
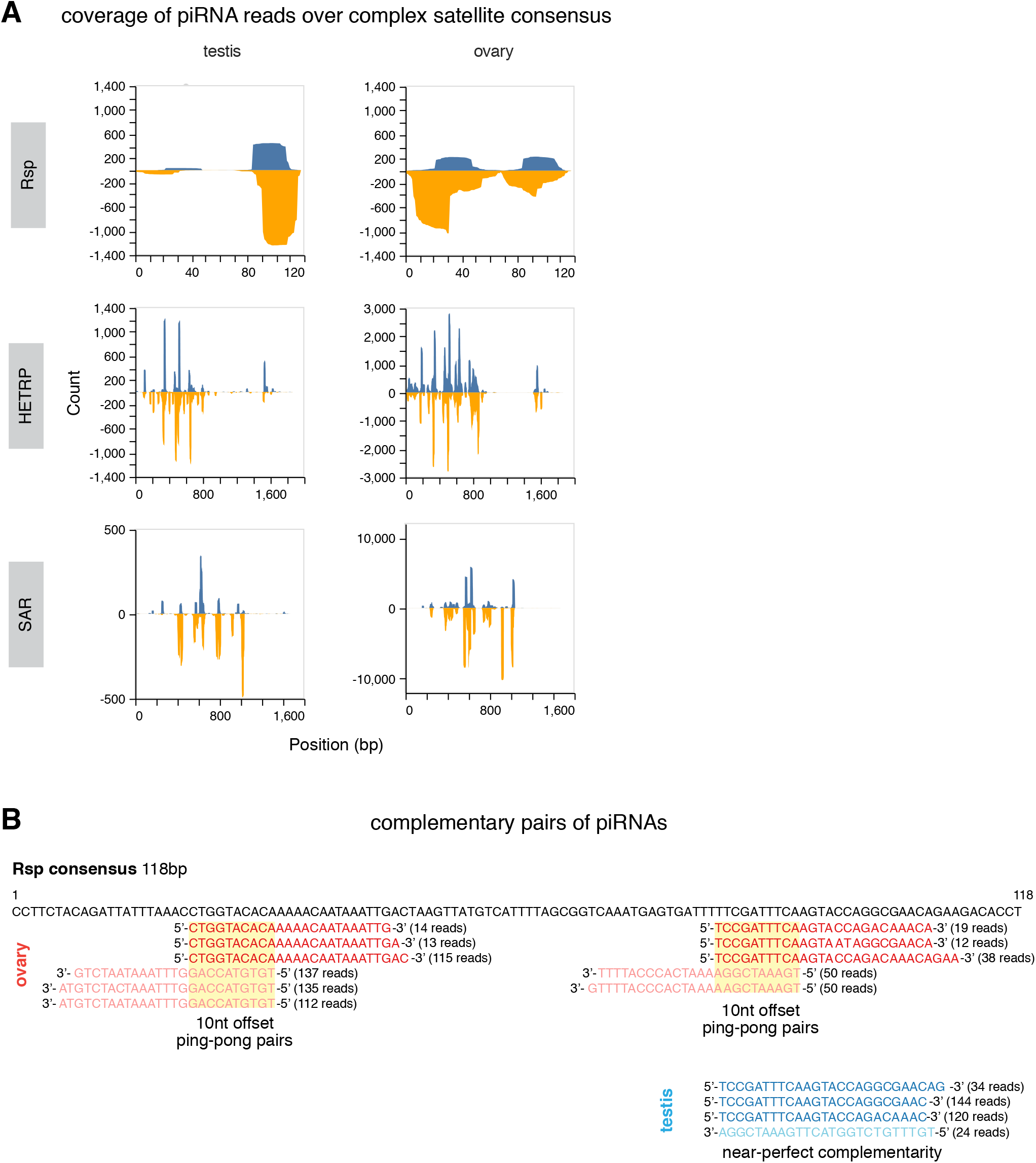
Coverage of piRNAs over consensus sequences of complex satellites and examples of complementary *Rsp*-mapping piRNA pairs in two sexes. Related to Figure 1. (A) Coverage plots of piRNAs over *Rsp* (top), *HETRP* (middle) and *SAR* (bottom), in testis (left) and ovary (right). (B) Examples of complementary pairs of *Rsp*-mapping piRNAs. Note that in ovary (red) they show an enrichment for 10-nt overlap, i.e., ping-pong signature, but in testis (blue) they show near perfect-complementarity with no evidence for ping-pong signature.

**Figure S3.**
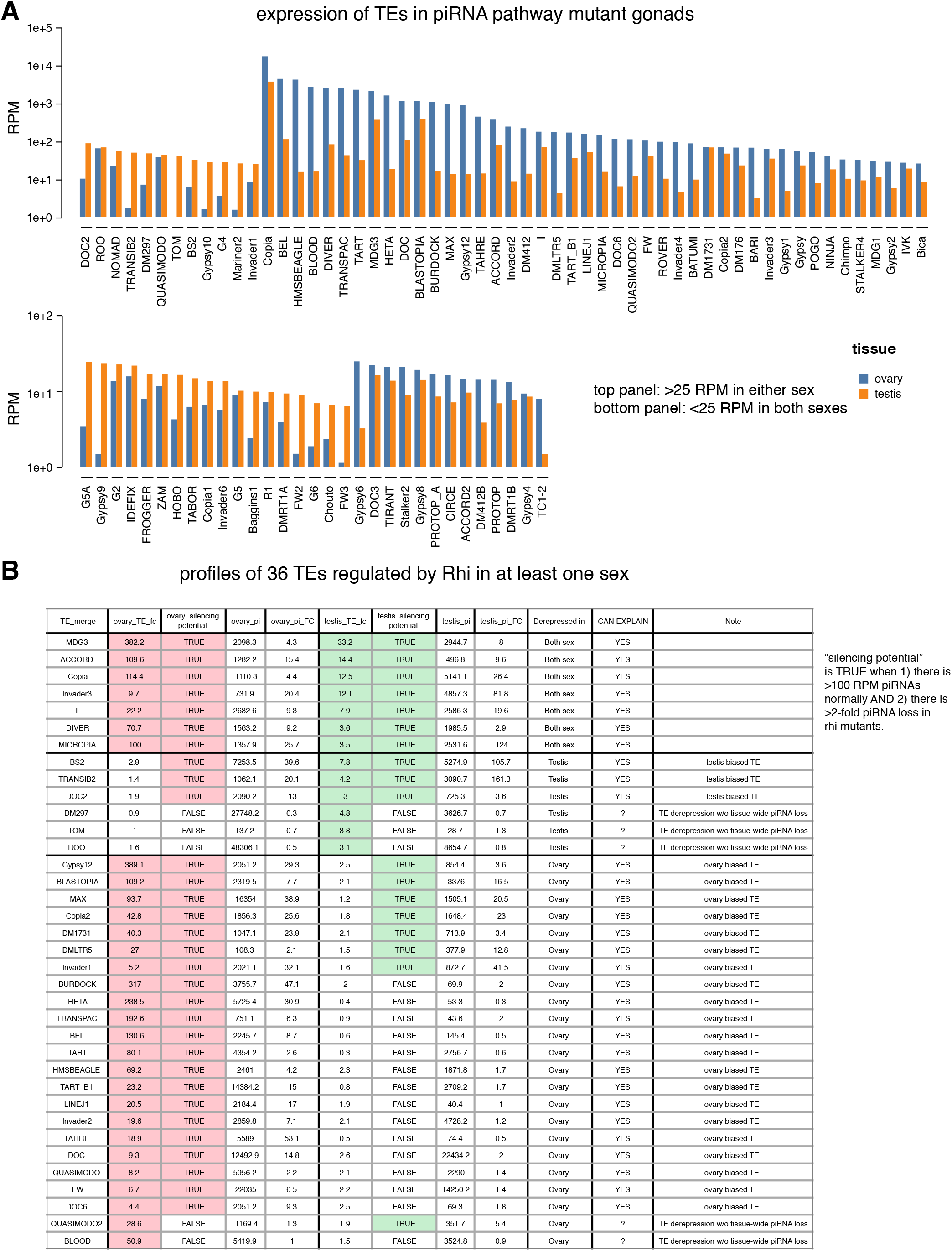
TE levels in piRNA pathway mutants and curation of TEs regulated by Rhi in at least one sex. Related to Figure 2,3. (A) Bar graphs showing TE levels in piRNA pathway mutant (*rhi*) testes (orange) and ovaries (blue). TEs that have at least 25 RPM in either sex is shown at the top, with the rest at the bottom. (B) Table reporting manual curation of 36 confidently affected TE families by *rhi^-/-^*. Silencing potential is TRUE when there are normally >100 RPM antisense piRNAs and they show >2-fold reduction in *rhi* mutants. TEs are deemed de-repressed when having >3-fold up-regulation. Note a few unexpected cases where TE de-repression is not accompanied by piRNA loss, the ovary ones of which were described before (Klattenhoff et al., 2009).

**Figure S4.**
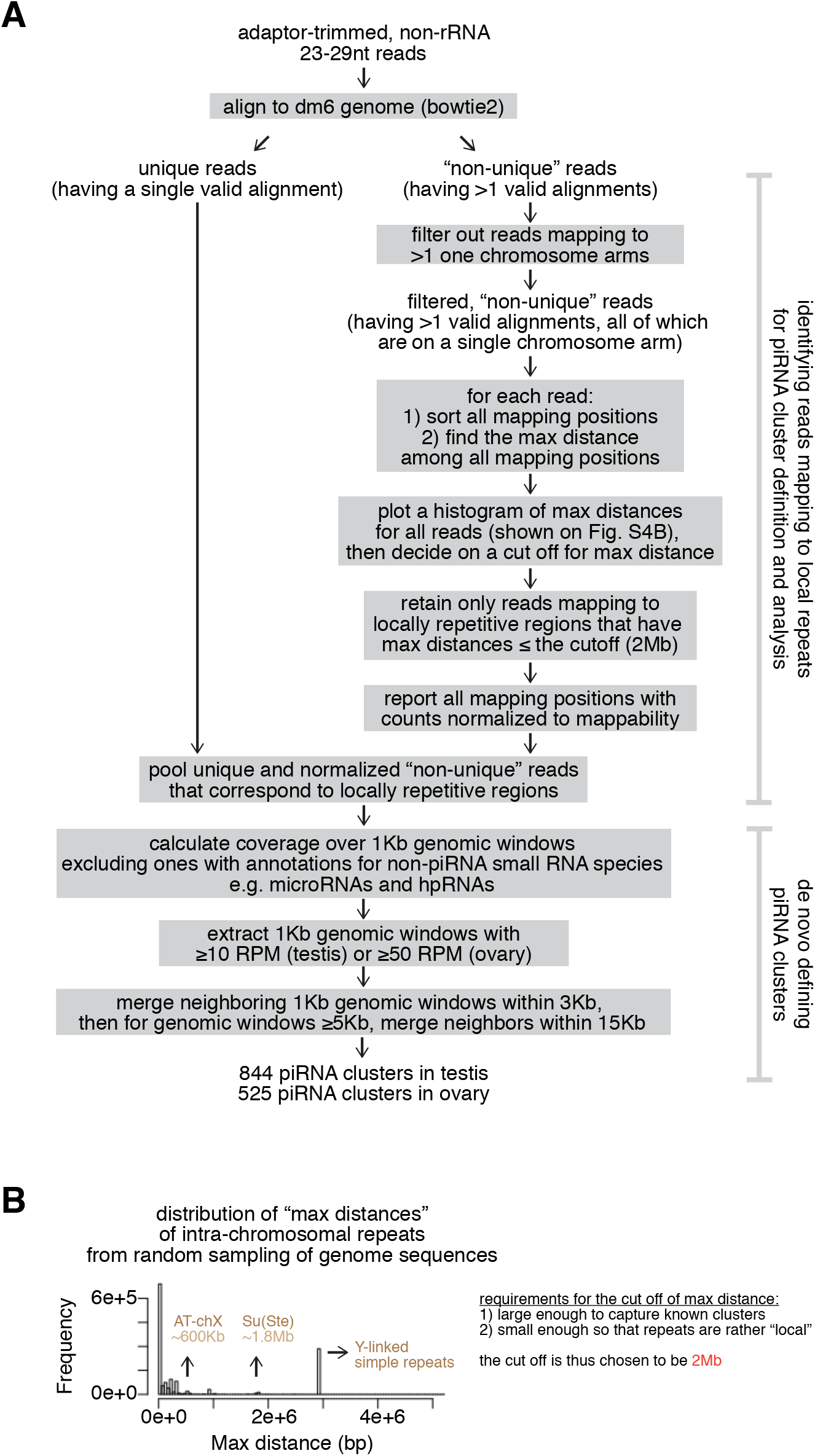
An algorithm that includes local repeats in piRNA cluster definition and analysis. Related to Figure 4. (A) Flow chart showing steps of the new algorithm that includes local repeats in piRNA cluster definition and analysis. See also methods. (B) Histogram showing the distribution of “max distances” defined in (A) to identify a meaningful cutoff (2Mb) for distinguishing local from non-local repeats. See also methods.

**Figure S5.**
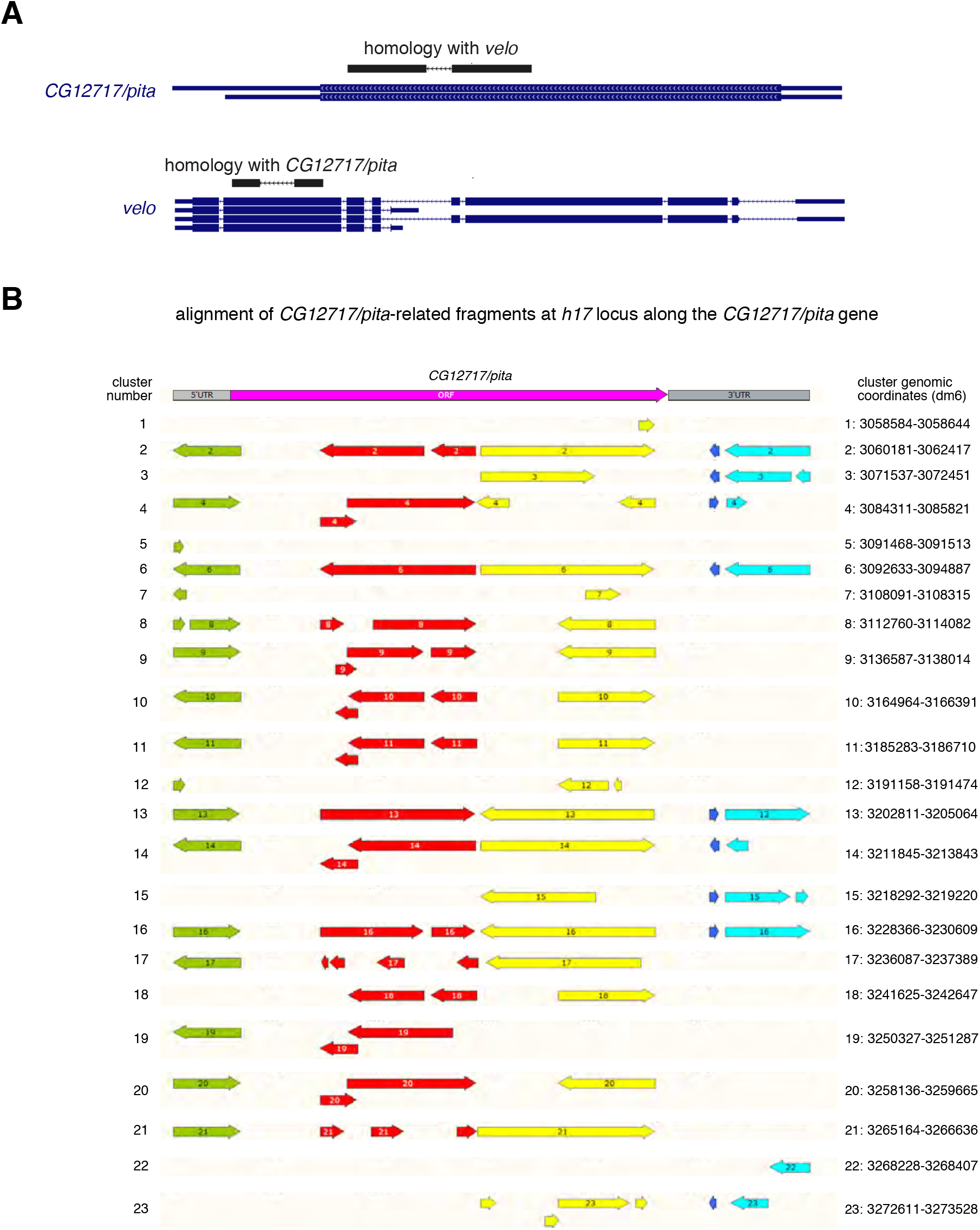
Characterization of *pita* homology in *D. mel*. Related to Figure 6,7. (A) Homology between two *D. melanogaster* paralogs: *velo* and *CG12717/pita*. The homologous regions are marked using BLAT and they share 75% nucleotide sequence identity. (B) Alignment of duplicated, partial copies of *CG12717* at *h17* on *D. melanogaster* Y to its *CG12717* gene (left), and their genomic coordinates (right). Note that there are two small regions of *CG12717* absent on Y. RNA *in situ* HCR was targeted against the ORF region unique to *CG12717* gene.

**Table S1. Genome-wide piRNA clusters in testis and ovary as well as major piRNA clusters defined in this study.**

## MATERIALS AND METHODS

### Fly stocks

Stocks and crosses were raised at 25 °C. The following stocks were used: *aub^QC42^* (BDSC4968), *aub^HN2^* (BDSC8517), *zuc^Df^* (BSDC3079), *spn-E^hls3987^* (BDSC24853) and *spn-E^1^* (BDSC3327) were obtained from Bloomington Drosophila Stock Center; *rhi^2^* and *rhi^KG^* were gifts of William Theurkauf; *zuc^HM27^* was a gift from Trudi Schüpbach; *nosP-GFP-Burdock* was a gift from Julius Brennecke. Heterozygous siblings were used as controls for all experiments.

### RNA *in situ* hybridization chain reaction (HCR)

A kit containing a DNA probe set, a DNA probe amplifier and hybridization, amplification and wash buffers were purchased from Molecular Instruments (molecularinstruments.org) for *CG12717* transcripts. To avoid targeting the *h17* region on Y, we designed probes against a ~400bp unique region present in *CG12717* on X but absent on Y chromosome. The *CG12717* probe set (unique identifier: 3916/E064) initiated B3 (Alexa546) amplifier. *In situ* HCR v3.0 (Choi et al., 2018) was performed according to manufacturer’s recommendations for generic samples in solution.

### Image acquisition and analysis

Confocal images were acquired with Zeiss LSM 800 using a 63x oil immersion objective (NA=1.4) and processed using Fiji (Schindelin et al., 2012). Single focal planes were shown in all images, where dotted outlines were drawn for illustration purposes.

### RNA-seq

RNA was extracted from 160-200 pairs of dissected testes of *aub^QC42/HN2^, spn-E^1/hls3987^, zuc^HM27/Df^* and respective heterozygous sibling controls in TRIzol (Invitrogen). PolyA+ selection was done using NEBNext Poly(A) mRNA Magnetic Isolation Module (NEB E7490), followed by strandspecific library prep with NEBNext Ultra Directional RNA Library Prep Kit for Illumina (NEB E7760) according to manufacturer’s instructions. Libraries were sequenced on Illumina HiSeq 2500 yielding 11-17 million 50bp single-end reads. PolyA-selected RNA-seq of *rhi* mutants and controls were downloaded from NCBI SRA (see the accompanying manuscript for testis and GSE126578 for ovary, 2 biological replicates per sex per genotype).

### RNA-seq analysis

To quantify expression levels of protein-encoding genes across different piRNA pathway mutants (*aub, zuc* and *spn-E*), we used kallisto 0.46.1 (Bray et al., 2016). Three heterozygous controls were pooled as triplicates of controls to be analyzed against duplicates of each of the three piRNA pathway mutants. Transcript-level quantification was pooled to obtain gene-level quantification. Differential gene expression was done with DESeq2 (Love et al., 2014). Expression of *CG12717* and *veloren* in ovary and testis from modENCODE (Brown et al., 2014) was extracted from FlyBase (Thurmond et al., 2019).

For analysis of TE expression and TE fold-change in piRNA pathway mutants of both sexes, *rhi* mutants were used where piRNA production from germline-specific dual-strand clusters was abolished. Reads mapped to rRNA were discarded using bowtie 1.2.2 allowing 3 mismatches. Reads were then mapped to TE consensus from RepBase17.08 using bowtie 1.2.2 with -v 3 -k 1 and normalized to the total number of reads mapped to dm6 genome. For simplicity, reads mapped to LTR and internal sequences were merged for each LTR TE given their well correlative behaviors. Only TEs that have ≥5 RPM expression in piRNA pathway mutants of either sex were kept for the analysis (n=87). A pseudo-count of 1 was added before calculating TE fold-change in piRNA pathway mutants.

### Identification of TEs regulated by *rhi*

To identify a set of TEs regulated by *rhi* in at least one sex, we looked for TEs that have at least 100 RPM in *rhi* mutant ovaries or at least 25 RPM in *rhi* mutant testes. Next, we filter out TEs that show less than 3-fold de-repression in both sexes. From the initial 87 TEs defined above, these led to a total of 36 TEs regulated by *rhi* in at least one sex shown in Figure 2E and Figure 3A. See Figure S3B for detailed profiles of these 36 TEs.

### piRNA-seq

RNA extraction was done as above for RNA-seq. 18-30nt small RNAs were purified by PAGE (15% polyacrylamide gel) from ~1μg total RNA. Purified small RNA was subject to library prep using NEBNext Multiplex Small RNA Sample Prep Set for Illumina (NEB E7330) according to manufacturer’s instructions. Adaptor-ligated, reverse-transcribed, PCR-amplified samples were purified again by PAGE (6% polyacrylamide gel). Two biological replicates per genotype were sequenced on Illumina HiSeq 2500 yielding 15-20 million 50bp single-end reads.

### piRNA-seq analysis of TEs, complex satellites and genes

To isolate piRNAs, adaptor-trimmed total small RNAs were size-selected for 23-29nt (cutadapt 2.5) and those mapped to rRNA, miRNA, snRNA, snoRNA and tRNA were discarded (bowtie 1.2.2 with -v 3). piRNAs were first mapped to RepBase17.08 to obtain the portion mapping to TEs and complex satellites; the rest was then mapped to gene sequences derived from the gtf file downloaded from Ensembl (BDGP6.28.99) (Yates et al., 2019); reads unmapped to repeats and genes were then mapped to dm6 to infer the portion mapping to inter-genic regions, and the unmapped ones were listed under “others” category. A pipeline is also drawn in Figure S1. For TE-antisense piRNA analysis, piRNA reads were mapped, normalized and processed as done for polyA+ RNA-seq (see above). For complex satellite-mapping small RNAs, we plotted size distribution, analyzed nucleotide bias at position 1 and calculated coverage along consensus sequences using bedtools v2.28.0. Ping-pong signature analysis (i.e., 5’-to-5’ distances between complementary piRNA pairs) was done with custom scripts. Ping-pong z-score was calculated using 1-9nt and 11-23nt as background distribution for an enrichment of 10nt. For piRNAs antisense to protein-encoding genes of interest, we downloaded gene sequences from FlyBase (Thurmond et al., 2019) and mapped piRNAs to them using bowtie 1.2.2. For mRNA-derived sense piRNAs, we mapped piRNAs to genome and kept ones with unique mapping and zero mismatch (bowtie 1.2.2 with -v 0 -m 1) to the gene regions and orientations of interest.

### A pipeline tolerating local repeats for piRNA cluster analysis

We first separated rRNA-depleted 23-29nt small RNA reads that map to one unique location in the genome and others that have multiple mapping positions (“multi-mappers”). For all multimappers, we filtered out those who map to more than one chromosome arm, retaining only ones with multi-mapping positions on a single chromosome arm (“intra-chromosomal repeats”). Then, for each of the reads we kept as intra-chromosomal repeats, we calculated the maximum distance (“max distance”) of all mapping positions. In order to enforce the local requirement, we hoped to identify a cutoff distance for max distances, which is large enough to contain known piRNA loci but small enough to allow certain resolution of neighboring loci. To this end, we analyzed a pool of 50bp DNA fragments tiling the entire dm6 genome and plotted a histogram of max distances for all intra-chromosomal repeats (Figure S4B). This revealed a density of intra-chromosomal repeats having max distances smaller than ~500Kb, as well as four pronounced peaks with larger max distances. Sequence analysis uncovered the identities of these peaks: the peak with ~600Kb max distance corresponds to *AT-chX*, the peak with ~1.8Mb max distance represents *Su(Ste*), and the other two peaks mostly contain Y-specific simple repeats. We thus set a 2Mb tolerance threshold of max distances to allow local repeats in piRNA cluster analysis. In other words, we defined local repeats as repeats that have all copies contained within a window smaller than 2Mb and merged their normalized counts with unique sequences for piRNA cluster analysis. Alignment was done using bowtie2 to dm6 genome. To compare this new pipeline with other standard approaches (permitting only unique mappers or allowing all multi-mappers with randomly assigned locations), we calculated the number of reads mapped to major piRNA clusters using different methods (Figure 4B). A summary of this pipeline is shown on Figure S4A.

### Definition of piRNA clusters

23-29nt small RNAs were mapped to dm6 genome using the above-mentioned pipeline tolerating local repeats and generated coverage profiles across 1Kb windows that tile the genome. 1Kb windows including highly expressed miRNA, snRNA, snoRNA, hpRNA or 7SL SRP RNA were excluded. 1Kb windows with low read-coverage (≤100bp) were also excluded. Then, 1Kb windows that produce at least certain amounts of piRNAs were extracted for cluster definition (≥10RPM for testis, ≥50RPM for ovary). Neighboring 1Kb widows within 3Kb were merged. If merged windows were ≥5Kb, they were merged again within 15Kb. This yields 844 piRNA clusters in testis and 525 piRNA clusters in ovary, after manual curation. Major piRNA clusters described before in ovaries (Brennecke et al., 2007; Mohn et al., 2014) were all recovered with similar resolution. To compare expression levels of major piRNA clusters between sexes, cluster boundaries were manually curated to guarantee identical regions being compared. piRNA clusters defined in this study for both sexes are listed in Table S1.

### TE content of piRNA clusters

TE annotation in dm6 genome was downloaded from UCSC Table Browser (Karolchik et al., 2004). piRNA cluster boundaries were defined as described above. For piRNA cluster of interest, the TE content is calculated as length contribution to the entire cluster length by individual TEs. TE contents add up to less than 100%, as TEs do not fill completely the cluster length.

### Sex bias of piRNA cluster TE content

Sex bias of individual TEs was first computed as log2 ratio of expression levels in piRNA pathway mutants (*rhi*) between sex (ovary over testis). Sex bias of piRNA cluster TE content was then computed as the cumulative sex bias of individual TEs inside the cluster, weighted by their length contribution to the cluster. Using all expressed TEs or only ones that show pronounced sex bias generated comparable results. To eliminate noise, we only used TEs that exhibit strong, ≥10-fold sexual difference in expression (n=24). An equation and an example are shown in Figure 5C.

### BLAT and BLAST analysis

To characterize the unannotated sequence between annotated repeats in piRNA clusters, interrepeat sequences were analyzed using BLAT on UCSC Genome browser (Kent, 2002). For example, an inter-TE sequence at *Hsp70B* locus was used to BLAT against dm6 genome, which revealed the homology with an exon of *nod* gene (Figure 6A). Homology between *CG12717* and *veloren* was done with both BLAT and BLAST, which yielded similar results. Characterization of *CG12717*-homologous sequences at *h17* locus (Figure S5B) was done by multiple sequence alignment with the Needle program (ebi.ac.uk/Tools/psa/emboss_needle/).

### Phylogenetic analysis

The longest transcripts of *veloren* and *CG12717* in *D. melanogaster* genome were used to BLAST against nucleotide collection with blastN program. Orthologs of these two genes in other *Drosophila* species were identified based on high nucleotide similarity and synteny. In all orthologs identified for both genes, we found the same flanking protein-encoding genes, confirming their ortholog identities. Occasionally, BLAST with *CG12717* revealed the *veloren* ortholog in that species as well; but only in *D. mauritiana, D. simulans* and *D. sechellia* genomes are there additional hits with high sequence homology to *CG12717*, other than the orthologous *CG12717* and *veloren*. These additional *CG12717*-related sequences are in some cases annotated as predicted genes, but all buried in TE-rich heterochromatin (close to centromere or in highly repetitive unassigned scaffolds). To examine the organization of *CG12717*-related sequences in *D. mauritiana* genome in detail, we ran BLAST using *D. mauritiana CG12717* gene against its genome (assembly: GCA_004382145.1), which revealed additional unannotated regions with high sequence similarity to *CG12717*. Those located on chrX and chr3 were drawn in Figure 7G. The instance where two adjacent *CG12717*-related sequences are arranged head-to-head on chrX is illustrated in Figure 7I, and the other three such instances are found in unassigned scaffolds. To uncover the identity of flanking unannotated sequences, we BLAST the 50Kb region encompassing *CG12717*-related sequences against TE consensus (RepBase17.08). The cladogram was drawn for illustration (Drosophila 12 Genomes Consortium, 2007).

### Analysis of testis small RNAs in non-*D. melanogaster* species

Testis small RNA libraries from non-*D. melanogaster* species was downloaded from NCBI SRA: *D. simulans* SRR7410589 (Lin et al., 2018) and *D. mauritiana* SRR7961897 (Kotov et al., 2019). Adaptor-trimmed reads were mapped to the orthologous *CG12717* gene, *D. simulans GD15918* and *D. mauritiana LOC117148327*, respectively (bowtie 1.2.2 with -v 3 -k 1). Coverage was plotted along the orthologous *CG12717* gene.

### Data visualization and statistical analysis

Most data visualization and statistical analysis were done in Python 3 via JupyterLab with the following software packages: numpy (Oliphant, 2015), pandas (McKinney, 2010) and altair (VanderPlas et al., 2018). The UCSC Genome Brower (Kent et al., 2002) and IGV (Robinson et al., 2011; Thorvaldsdóttir et al., 2013) were used to explore sequencing data and to prepare browser track panels shown.

### Data and code availability

Sequencing data will be uploaded to NCBI SRA.

